# Very rapid flow cytometric assessment of antimicrobial susceptibility during the apparent lag phase of microbial (re)growth

**DOI:** 10.1101/480392

**Authors:** Srijan Jindal, Harish Thampy, Philip J. Day, Douglas B. Kell

## Abstract

Cells of *E. coli* were grown in LB medium, taken from a stationary phase of 2-4h, and reinoculated into fresh media at a concentration (10^5^.mL^-1^ or lower) characteristic of bacteriuria. Flow cytometry was used to assess how quickly we could detect changes in cell size, number, membrane energisation (using a carbocyanine dye) and DNA distribution. It turned out that while the lag phase observable macroscopically via bulk OD measurements could be as long as 4h, the true lag phase could be less than 15-20 min, and was accompanied by many observable biochemical changes. Antibiotics to which the cells were sensitive affected these changes within 20 min of reinoculation, providing the possibility of a very rapid antibiotic susceptibility test, on a timescale compatible with a visit to a GP clinic. The strategy was applied successfully to genuine potential Urinary Tract Infection (UTI) samples taken from a doctor’s surgery. The methods developed could prove of considerable value in ensuring the correct prescription and thereby lowering the spread of antimicrobial resistance.

## Introduction

As is well known, there is a crisis of resistance to antimicrobials [1-8], caused in part by mis-prescribing. This mis-prescribing takes two forms: (i) potentially effective antibiotics are given when the infection is not bacterial, or (ii) the wrong (i.e. ineffective) antibiotics are given when it is. What would be desirable would be a very rapid means of knowing, even before a patient left a doctor’s surgery, that a particular antibiotic was indeed capable of killing or inhibiting the growth of the organism of interest. While genotypic (whole-genome-sequencing) methods hold out some promise for this [9-19], what is really desired is a phenotypic assay [20] that assesses the activity of anti-infectives in the sample itself [21-23]. However, since almost all antibiotics, whether bacteriostatic or bactericidal [24], indicate their efficacy or otherwise only when cells are attempting to replicate [25], it might be thought that this would be an unattainable goal simply because of the existence of a lag phase (but see below).

Urinary tract infections (“UTIs”) are a worldwide patient problem [22]. Other than in hospital-acquired infections [26], they are particularly common in females, with 1 in 2 women experiencing a UTI at some point in their life [27]. *Escherichia coli* is the most common causative pathogen of a UTI [27-30]. However, other Enterobacteriaceae such as *Proteus mirabilis*, *Klebsiella* spp. and *Pseudomonas aeruginosa*, and even Gram-positive cocci such as staphylococci and enterococci, may also be found [31, 32]. Misapplication and overuse of antibiotics in primary care is a major source of antimicrobial resistance [26, 32, 33], so it is important that the correct antibiotic is prescribed [34, 35]. Often prescribing none at all for asymptomatic UTIs is an adequate strategy [36].

We note, however, that *E. coli cells* in all conditions are highly heterogeneous [37], even if only because they are in different phases of the cell cycle [38], and in both ‘exponential’ and stationary phase contain a variety of chromosome numbers [39-44]. To discriminate them physiologically, and especially to relate them to culturability (a property of an individual), it is necessary to study them individually [45, 46], typically using flow cytometry [45, 47-56]. Flow cytometry has also been used to count microbes (and indeed white blood cells) for the purposes of assessing UTIs [57-61], but not in these cases for antibiotic susceptibility testing. Single cell morphological imaging has also been used, where in favourable cases antibiotic susceptibility can be detected in 15-30 minutes or less [62, 63].

### Flow cytometry and antimicrobial susceptibility

A number of workers have recognised that flow cytometry has the potential to detect very rapid <u>changes</u> in both cell numbers, morphology (by light scattering) and physiology (via the addition of particular fluorescent stains that report on some element(s) of biochemistry or physiology). Boye and colleagues could see effects of penicillin on flow cytograms within an hour of its addition to sensitive strains [47]. Similarly, Gant and colleagues [64] used forward and side scattering, and noticed antibiotic-dependent effects on the profiles after 3h, but did not measure absolute counts. Later studies [65, 66] used the negatively charged dye bis-(1,3-dibutyl-barbituric acid) trimethine oxonol (DiBAC4(3)), which increases its binding and hence fluorescence upon loss of membrane energisation (that decreases the activity of efflux pumps such as acrAB/tolC [67-69]); they could detect susceptibility to penicillin and gentamicin in 2-5h. Using a similar assay, Senyurek and colleagues [70] could detect it within 90 min. Other workers have used a variety of probes, but evaluation was after a much longer period, e.g. 24h [71]. Álvarez-Barrientos and colleagues [72] give an excellent review of work up to 2000, with some reports (e.g. [73]) of detection of flow cytometrically observable changes in morphology (light scattering) at 30 min exposure to antibiotic. Flow cytometry has also been used to detect bacteriuria, although the numbers found seemed not to correlate well with CFUs [74]. Most so-called ‘live/dead’ kits rely on the loss of membrane integrity to detect the permeability of DNA stains, but many effective antibiotics have little effect on this in the short term, and such kits do not assess proliferation [75]. Because different probes and different antibiotics have different effects (and with different kinetics) on membrane integrity, we decided that the best strategy would be to look at the ability of antibiotics to inhibit proliferation directly, distinguishing bacteria from non-living scattering material via the use of a positively charged dye that energised living cells accumulate. Rhodamine 123 is a very popular dye of this type, but without extra chemical treatments that would inhibit proliferation is effective only in Gram-positive organisms [76, 77]. However, the positively charged carbocyanine dye 3,3′-dipropylthiadicarbocyanine iodide (di-S-C3(5)) [78, 79] seems to bind to and/or be accumulated by both Gram-positive and -negative bacteria, and provides a convenient means of detecting them.

### The ‘lag’ phase

A classical activity of general and laboratory microbiology involves the inoculation of a liquid nutrient broth with cells taken from a non-growing state, whether this be from long-term storage (typically in agar) or using cells that have been grown to stationary phase [80], more or less recently, in another liquid batch culture. The result of this is that the cells will, in time, typically increase in number and/or biomass, often exponentially, but that this is preceded by a ‘lag phase’ (that may be subdivided [81]) before any such increases. The length of the lag phase depends on various factors, including the nature of the nutrient media before and after inoculation, the inoculum density, pH, temperature, and the period of the previous stationary phase for that cell [82-88]. It is usually estimated (and indeed defined) by extrapolating to its starting ordinate value a line on a plot of the logarithm of cell number, cfu or biomass against time (e.g. [82, 89-94]). However, because of the different (and generally lower) sensitivity of bulk optical estimates of biomass [82, 95, 96] (and see later, Figure 1), only the first two of these are normally considered to estimate the ‘true’ lag phase.

**Figure 1.**
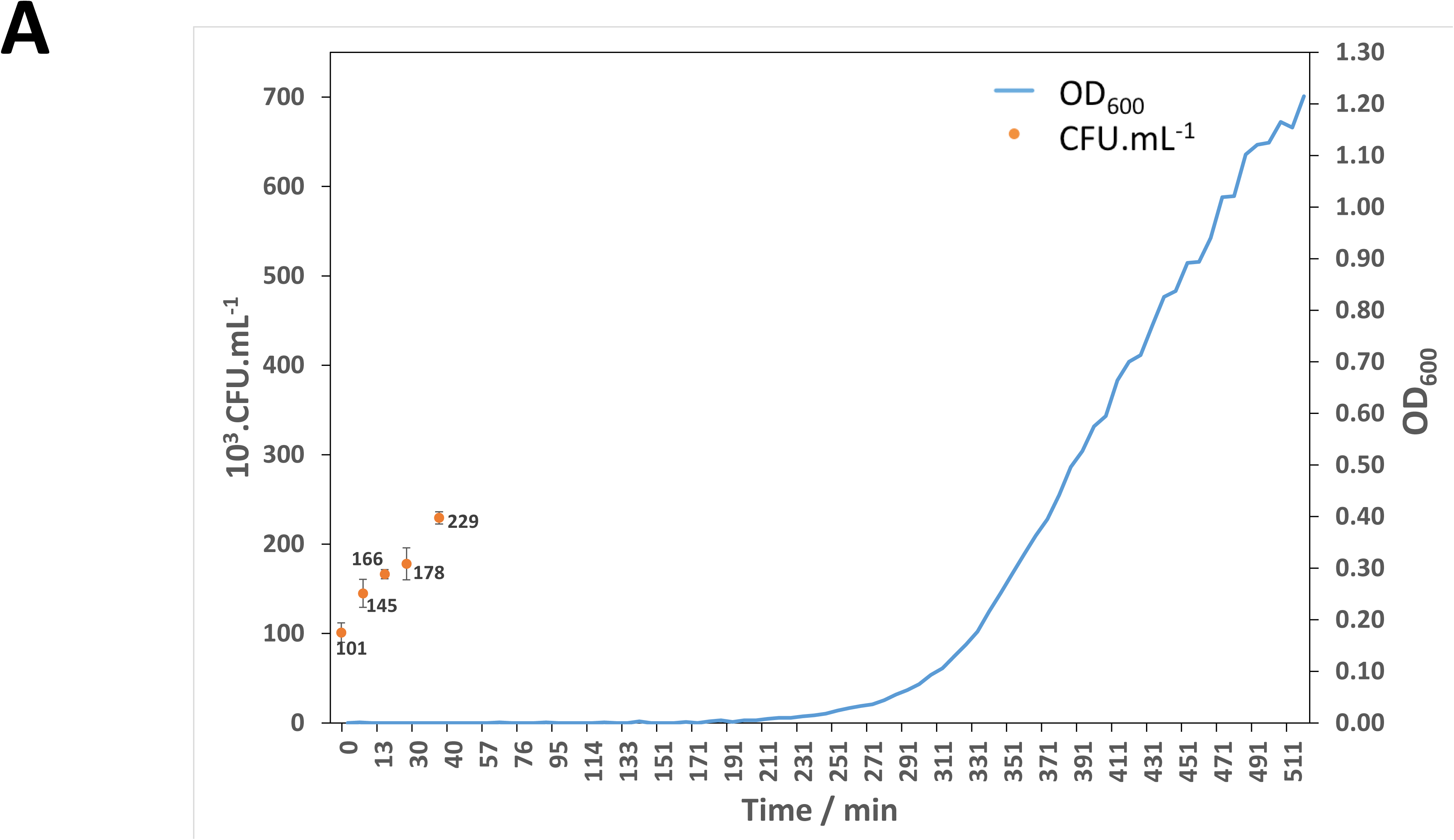
True and apparent lag phases during microbial regrowth. The strains indicated were grown in Lysogeny Broth and inoculated into Terrific Broth after 4 h in stationary phase to ca 10^5^ cells.mL^-1^. OD was measured quasi-continuously in An Omega plate reader spectrophotometer (BMG Labtech, UK), while CFU were measured conventionally on agar plates containing nutrient agar medium solidified with 1.5% agar. The lag phase observed via counting CFUs is about 30 min while bulk OD measurements show a lag phase of some 230 minutes (~4 hours).

With some important exceptions (e.g. [84, 85, 95, 97-99], the lag phase has been relatively little studied at a molecular level. From an applied point of view, however, at least two influences on it are considered desirable. Thus, a food microbiologist might wish to maximise the lag phase (potentially indefinitely) (e.g. [100]). By contrast, there are circumstances, as here, and not least in clinical microbiology, where it is desirable to be able to measure microbial growth/culturability, and its phenotypic sensitivity or otherwise to candidate anti-infective agents, in as short a time as possible. This necessarily involves minimising the length of the lag phase, and is the focus of the present studies.

There is evidence that the time before measurable biochemical changes occur during lag phases can be very small when inoculation is into rich medium [95, 98, 99]. Thus, Rolfe and colleagues [98] used Lysogeny Broth (LB) (and *S. enterica*), where lag phase or regrowth – as measured by changes in the transcriptome – initiated **within 4 min** (the earliest time point measured). The timescale in the plots of Madar and colleagues [95] does not admit quite such precise deconvolution, but responses in M9 with casamino acids (referred to as ‘immediate’) are consistent with a period of **less than 10 min**. Hong and colleagues recently detected such changes in under 30 min using stimulated Raman imaging [101], Yu and colleagues could do so with video microscopy [102], and Schoepp *et al*. [103] used molecular detection of suitable transcripts. In view of the above, and recognising that bacteria in UTIs may actually be growing (albeit slowly) and not in a ‘stationary’ phase, we decided to assess the ability of quantitative flow cytometry to determine bacterial cell numbers, and the effects of antibiotics thereon, on as rapid a timescale as possible. The present findings show that it is indeed possible to discriminate antibiotic-susceptible and – resistant strains in under 30 mins at levels (10^4-5^.mL^-1^) characteristic of bacteriuria [104, 105]. This opens up the possibility of ensuring that a correct prescription is given between the presentation of a sample in a doctor’s surgery and the acquisition in a dispensary of the correct antibiotic. A preprint has been lodged at bioRxiv.

## Materials and methods

### Microbial strains

*E. coli* MG1655 and a series of sensitive and resistant strains were taken from the laboratory collection of Prof R. Goodacre [106, 107].

### Culture

E. coli strains were grown from inocula of appropriate concentrations in conical flasks using Lysogeny Broth to an optical density (600nm) of 1.5 – 2, representing stationary phase in this medium. They were held in stationary phase for 2-4h before being inoculated at concentration of 10^5^ cells.mL^-1^ (or as noted) into Terrific Broth [108]. We did not here study cells held in a long stationary phase [80, 83] (exceeding 3d).

### Assessment of growth by bulk OD measurements

Bulk OD measurements were performed in 96-well plates and read at 600nm as per the manufacturer’s instructions in an Omega plate reader spectrophotometer (BMG Labtech, UK) instrument. The ‘background’ due to scattering from the plates, etc., was not subtracted.

### Flow cytometry

Initial studies used a Sony SH-800 instrument, but all studies reported here used an Intellicyt^®^ iQue screener PLUS. This instrument is based in significant measure on developments by Sklar and colleagues (e.g. [109-111]), and uses segmented flow [112] to sample from 96- or 384-well U- or V-bottom plates prior to their analysis. The iQue Plus contains three excitation sources (405nm, 488nm, 640nm) and 7 fixed filter detectors (with a midpoint/range in nm of 445/45, 530/30, 572/28, 585/40, 615/24, 675/30, 780/60, giving 13 fluorescence channels) whose outputs are stored as both ‘height’ and area, using the FCS3.0 data file standard [113]. Forward and side scatter are obtained from the 488nm excitation source. Detection channels are referred to by the laser used (405nm violet VL, 488 nm blue BL, and 640 nm red RL) and the detector number in order of possible detectors with a longer wavelength. Thus RL1, as used for detecting di-S-C3(5), implies the red laser and the 675/30 detector. Data are collected from all channels, using a dynamic range of 7 logs. Many parameters may be used to vary the precise performance of the instrument. Those we found material to provide the best reproducibility and to minimise carryover, and their selected values, are as follows: Automatic prime – 60 secs (in Qsol buffer); Pre-plate shake – 15 s and 1500rpm; Sip time – 2 s (actual sample uptake); Additional sip time – 0.5 s (the gap between sips); Pump speed – 29 rpm (1.5 µL.s^-1^ sample uptake); Plate model – U-bottom well plate (for 96 well plates); Mid plate cleanup – After every well (4 washes; 0.5 s each in Qsol buffer); Inter-well shake – 1500 rpm; after 6 wells, 4 sec in Qsol buffer; Flush and Clean – 30 sec with Decon and Clean buffers followed by 60 sec with deionised water. The Forecyt ^™^ software supplied with the instrument may be used to gate and display all the analyses post hoc. It, and the FlowJo software, were used in the preparation of the cytograms shown. Where used, di-S-C3(5) was present at a final concentration of 3 µM; its analysis used excitation at 640 nm and detection at 675±15 nm, the fluorescence channel being referred to as RL1. For DNA analyses, cells were fixed by injection into ice-cold ethanol (final concentration 70%), washed twice by centrifugation in 0.1 M-Tris/HCl buffer, pH 7.4, before resuspension in the same buffer containing mithramycin (50 µg mL^-1^) and ethidium bromide (25 µg mL^-1^), MgCl_2_, (25 mM) and NaCl (100 mM) [47]. Under these circumstances, the excitation energy absorbed by mithramycin (excitation 405 or 488nm) is transferred to the ethidium bromide, providing a large Stokes shift (emission at 572, 585 or 615 nm; we chose 572 nm as it provided the best signal) and high selectivity for DNA (as mithramycin does not bind to RNA). All the solutions and media used were filtered through 0.2 µm filter.

### UTI samples

Following ethics approval from the University of Manchester and the obtaining of signed consent forms, patients attending the Firsway clinic with suspected UTI were offered the opportunity to have their urine samples analysed by our method as well as the reference method used in a centralised pathology laboratory. Samples were taken at various times during the day, kept at 4°C, delivered to the Manchester laboratory by taxi, plated out (LB agar containing as appropriate the stated antibiotics at 3 times normal MIC) to assess microbial numbers and antibiotic sensitivity, the remaining sample kept again at 4°C, and analysed flow cytometrically within 18h. For flow cytometric assessment, cells were diluted into 37°C Terrific Broth containing 3 µM diS-C3(5) plus any appropriate antibiotic, and assayed as above. For other experiments (not shown) cells were filtered (0.45 µm) and diluted as appropriate into warmed Terrific Broth. No significant differences were discernible between the two methods.

### Reagents

All reagents were of analytical grade where available. Flow cytometric dyes were obtained from Sigma-Aldrich.

## Results

### Initial assessment of regrowth by bulk light scattering measurements in 96-well plates

Figure 1 shows a typical lag phase from an inoculum of 10^5^ cells.mL^-1^ that had spent 4h in stationary phase when inoculated into Terrific Broth [108] as observed by bulk OD measurements. For strain MG1655 the lag phase amounted to some 230 min, while as expected it is lower (2.5-3h) for the more virulent clinical isolates (not shown). A rule of thumb states that an OD of 1 is approximately equal to 0.5 mg.mL^-1^ dry weight bacteria or ~10^9^.cells.mL^-1^ for *E. coli*. Thus, the change in OD if 10^5^ cells.mL^-1^ increase their number by 50% is ~0.00015, which is immeasurably small in this instrument. Given the noise in the system (probably mainly due to fluctuations in the incident light intensity), it is reasonable that we might, in this system, detect changes in OD of 0.01 (~10^7^ cells.mL^-1^), which requires a 100-fold increase in cell number over the inoculum (~7 doublings). With a true lag phase of 10-15 min, and a doubling time of 20 min, this is indeed roughly what can be observed (Figure 1)(see also [114-116]). When samples were taken from the same strain and plated out to estimate proliferation by CFU, the results were indeed equally consistent with those at the longer times (Figure 1).

### Flow cytometric assessment of cells and cell proliferation

Figure 2A shows a typical set of traces of multiple wells from the Intellicyt iQue, each containing an inoculum of 10^5^ cells.mL^-1^. Each analysis is of 3 µL (taking 2s), and the good reproducibility is evident, especially in the inset stacked plot. Figure 2B rear trace shows the cytograms of a bead cocktail displaying that the distribution in cell properties is significantly greater than that of beads, and its significant width thus is not due to any inadequacies in the detector. The quality of a ‘high-throughput’ (or indeed any other) assay is nowadays widely assessed using the Z’ statistic [117]. This is given, for an assay in which the sample’s readout exceeds that of the control, as

**Figure 2.**
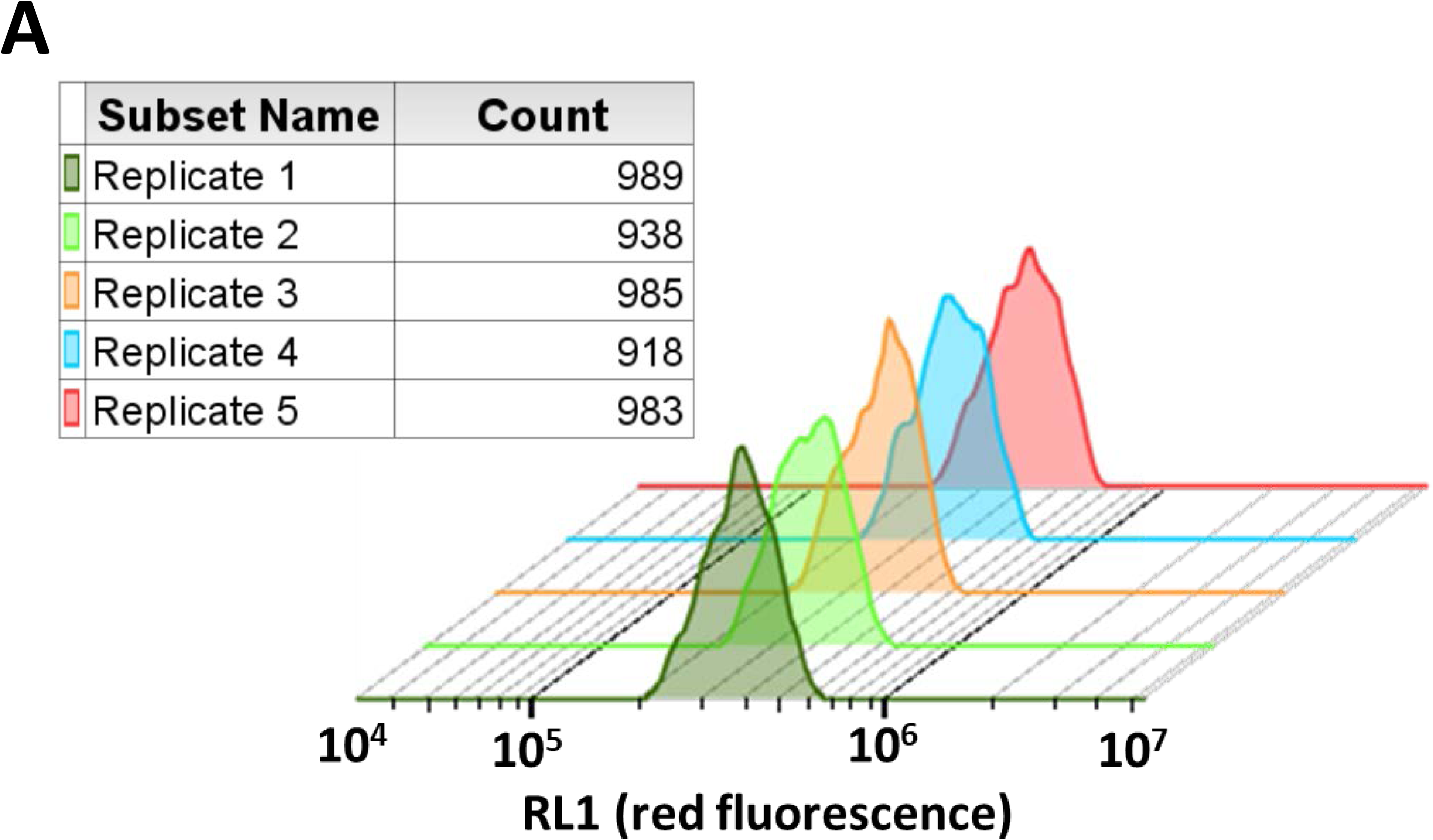
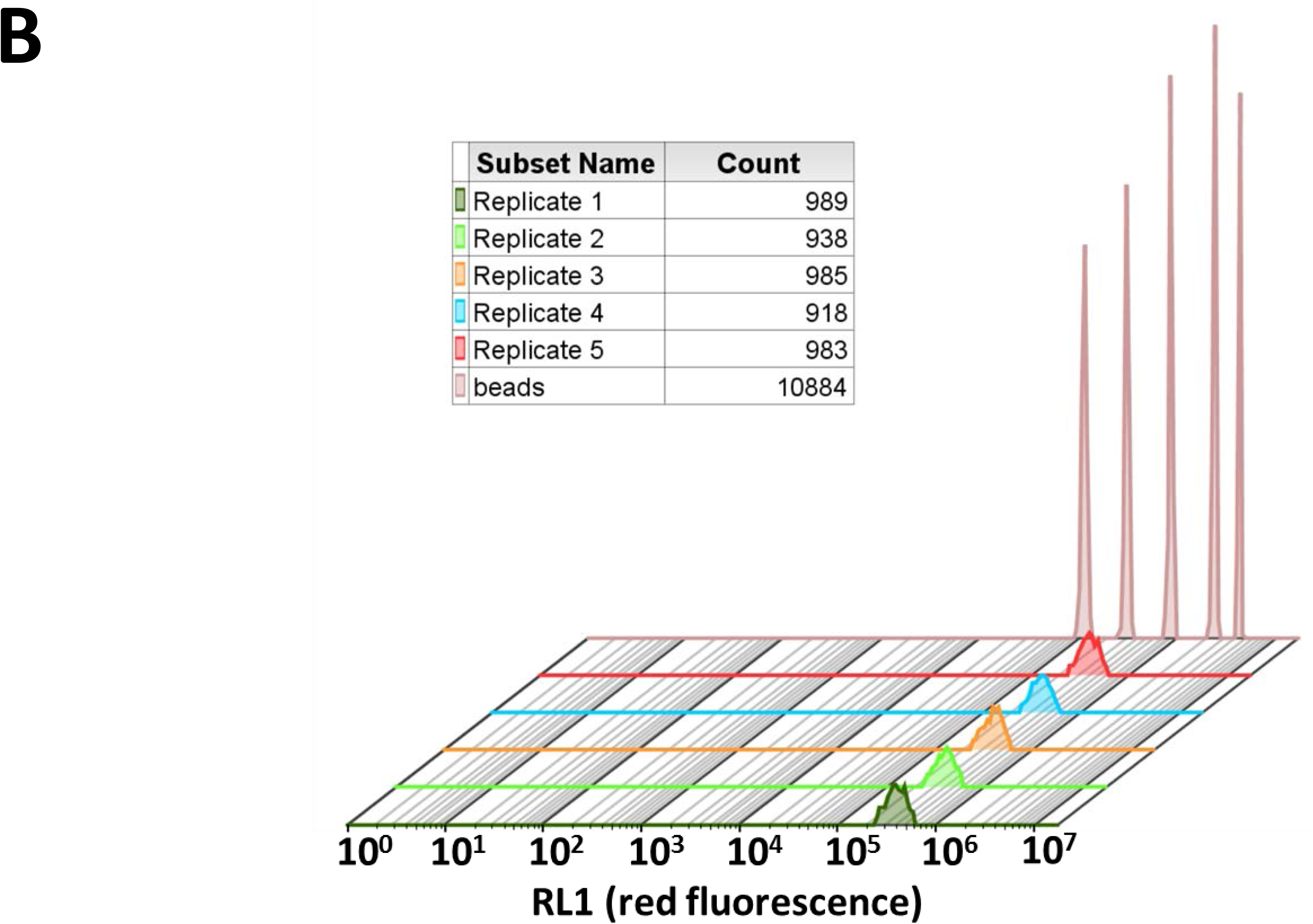
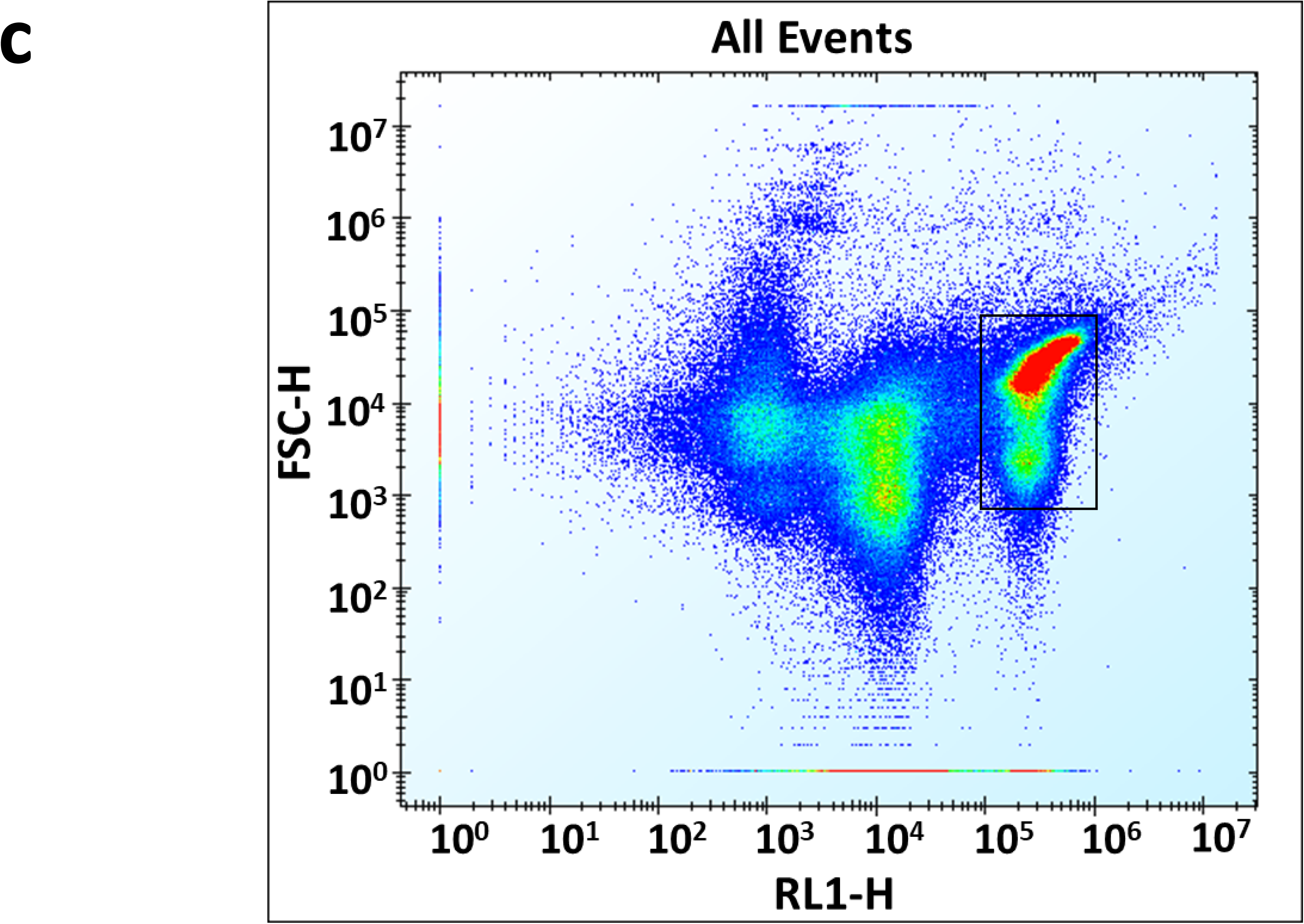
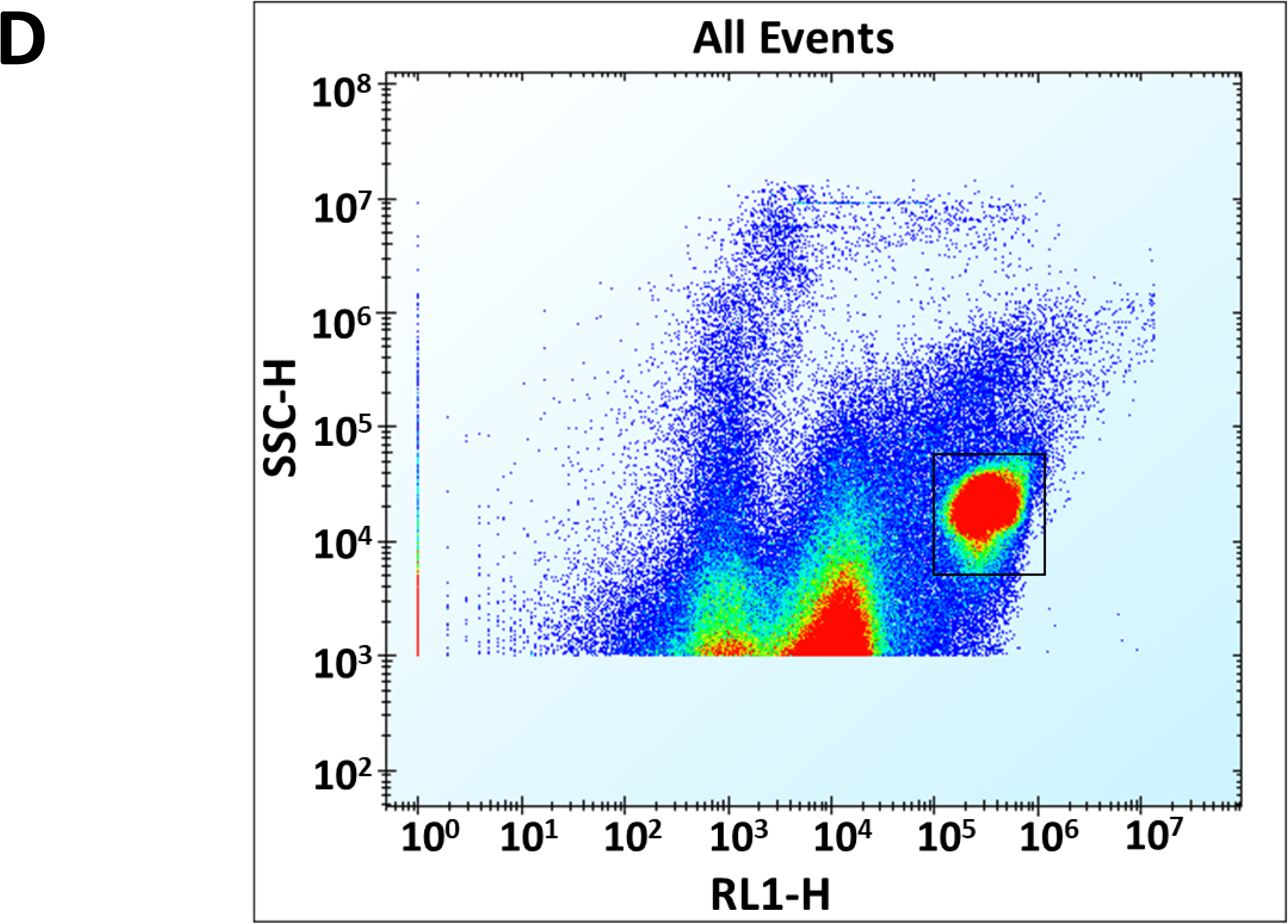
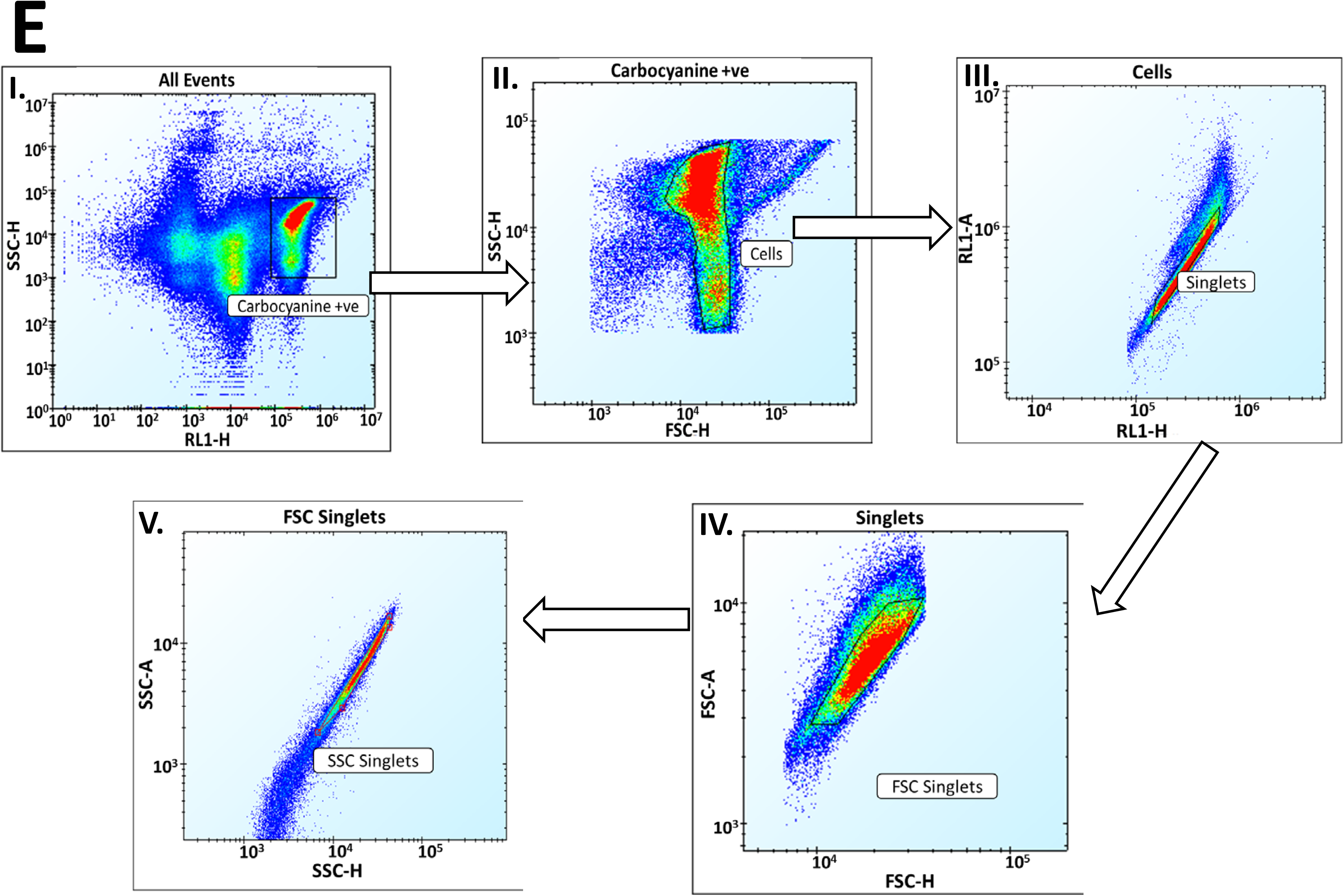
Cytograms of *E. coli* at a concentration of 10^5^ cells.mL^-1^ when incubated in 0.2 µm-filtered Terrific Broth containing 3µM Di-S-C3(5). The sample measured has a volume of 3µL and measurement takes place over 2 seconds. **A**. 1D histograms of RL1 fluorescence showing the reproducibility of the results. **B**. 1D histograms of RL1 fluorescence together with calibrating beads, showing that the breadth of the bacterial peaks is ‘real’ and not simply due to detector variability. **C** and **D**. Raw dot plots of the height of the forward scatter and of side scatter signals respectively vs RL1. Note that the *E. coli* cells appear above 10^5^ in RL1, the rest of the signals being due to very tiny unfiltered debris. **E**. Gating strategy (I-V) to show only the *E. coli* singlet cells.

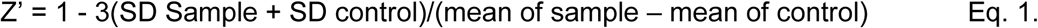

It is normally considered [117] that a Z’ factor exceeding 0.5 provides for a satisfactory assay.

Figures 2C and 2D show the full cytograms for forward scatter and side scatter, respectively vs RL1, illustrating the amount of small particulates remaining in Terrific Broth, despite extensive filtering. Consequently, we used a series of gates to assess solely the bacteria in our samples. These are shown in Figure 2E.

Figure 3 shows cytograms at various times after inoculation of the stationary phase (LB-grown) cells into Terrific Broth, along with labels of cell numbers within the regions of interest. These allow the assessment of the Z’ values as per equation 1. From Fig 3B it may be observed that Z’ > 0.5 from as early as 20 min, this then representing the earliest that we can robustly detect <u>proliferation</u>. Changes in cell constitution as judged by light scatter can, however, be detected from the earliest time point (5 min, Fig 3A top left). It is noteworthy that the proliferation (as measured by the increase in cell numbers on the ordinate) is parallelled, at least initially, by an increase in uptake of the carbocyanine dye (on the abscissa); as the cells ‘wake up’ they become increasingly energised, until they settle down (also observed via side scatter). For a lower concentration of starting inoculum (5×10^4^ cells.mL^-1^), the Z’ > 0.5 from 25 min as shown in figure 3C.

**Figure 3.**
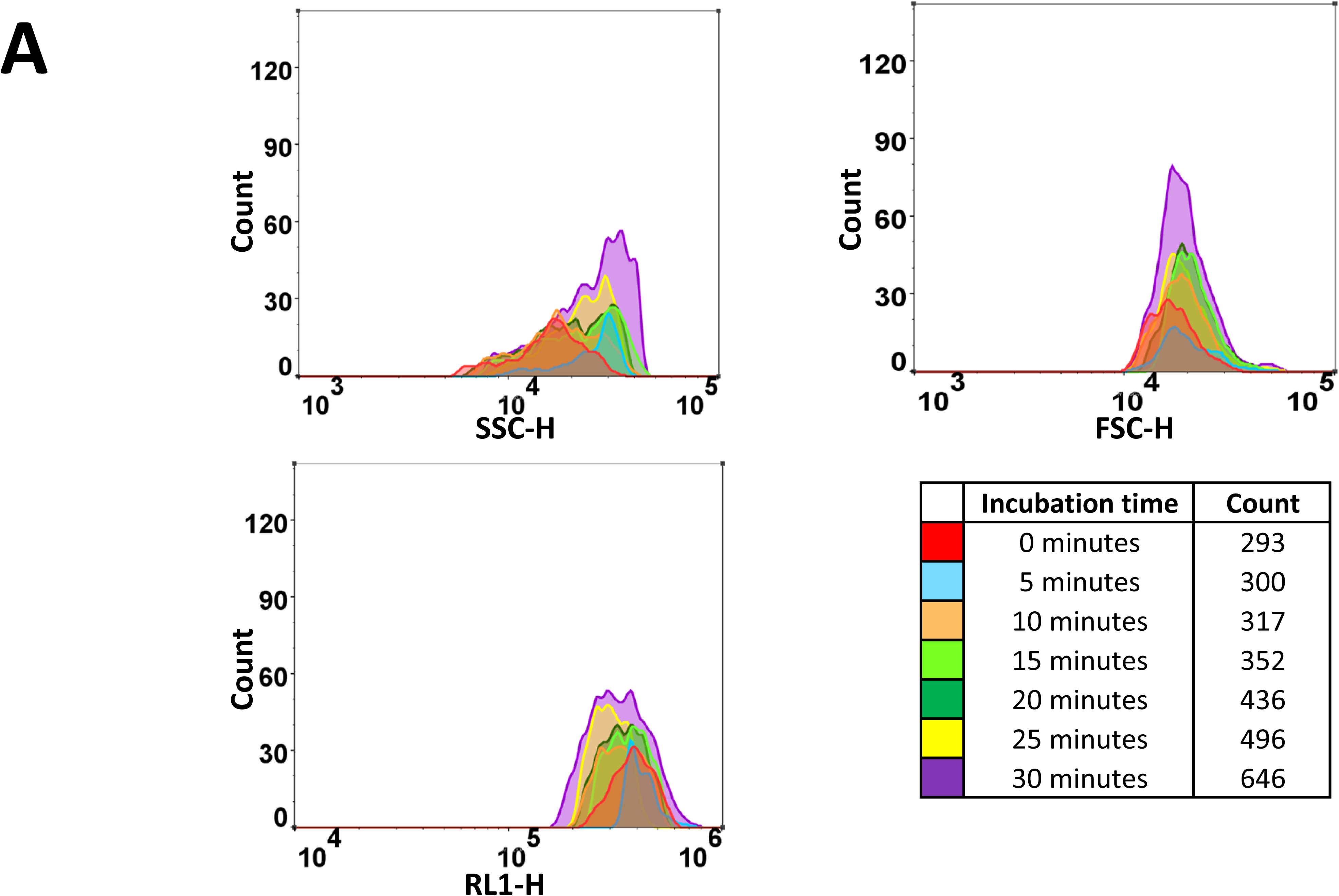
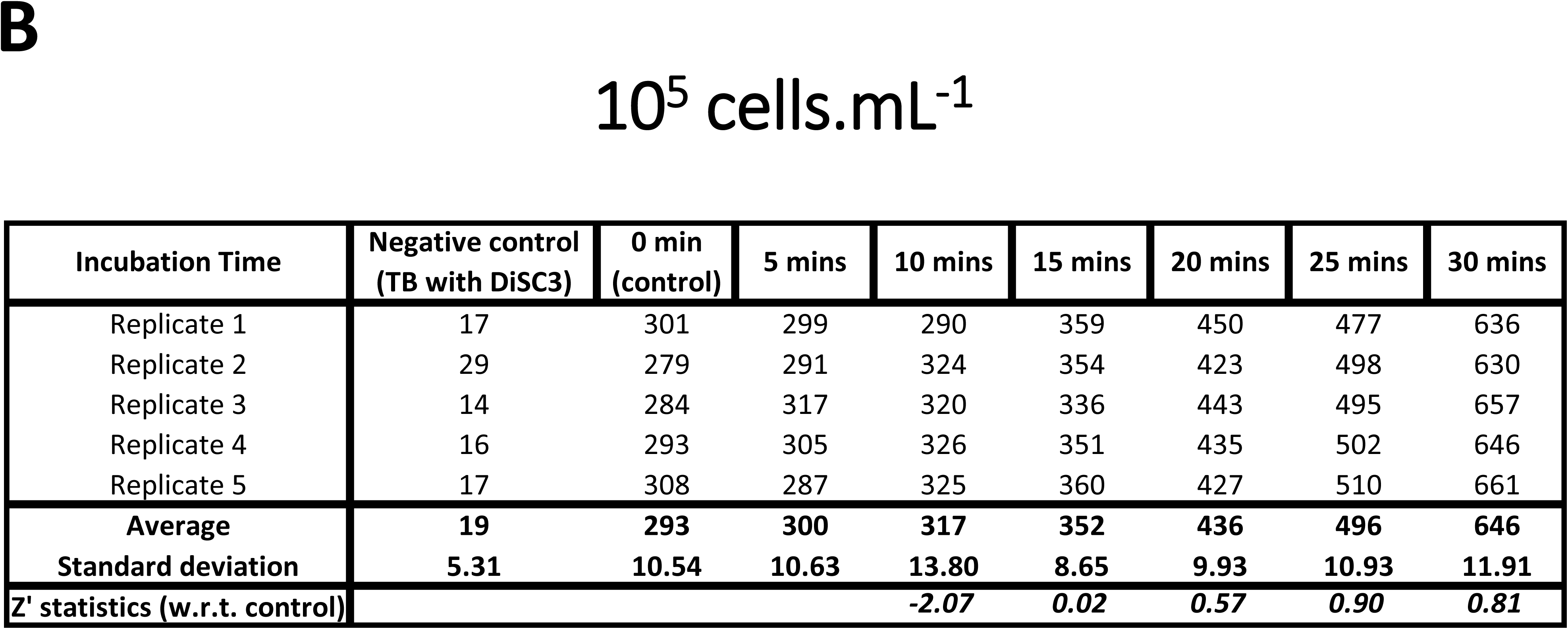
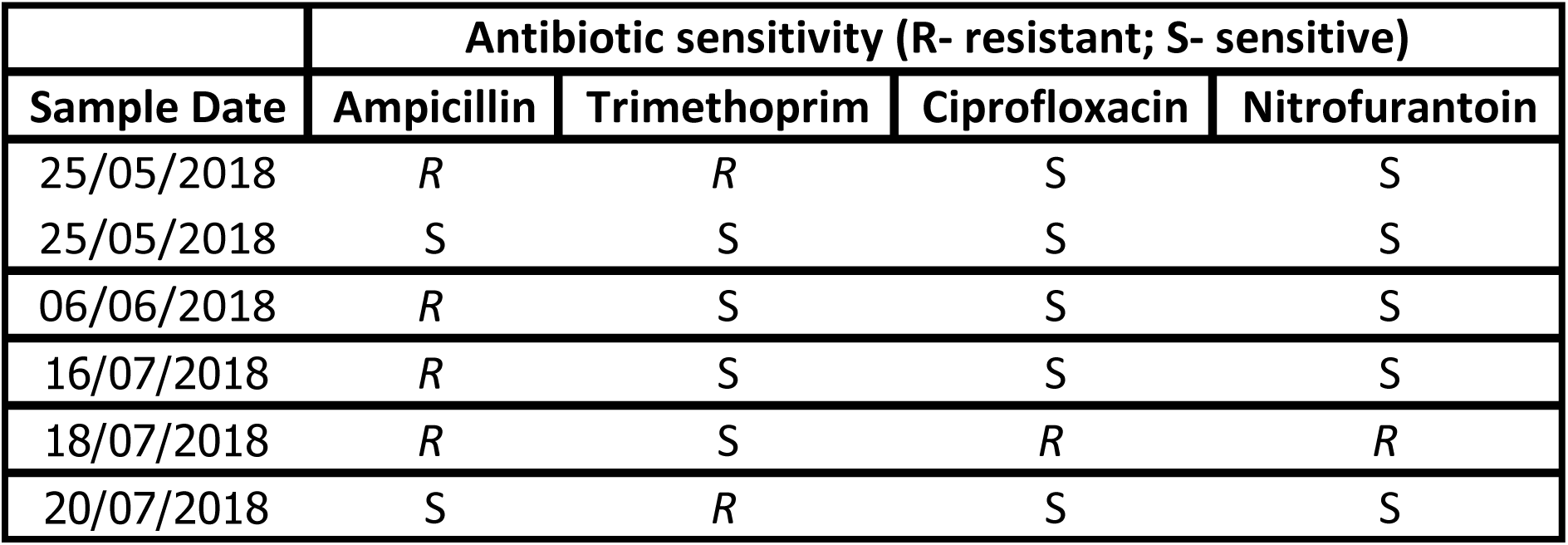
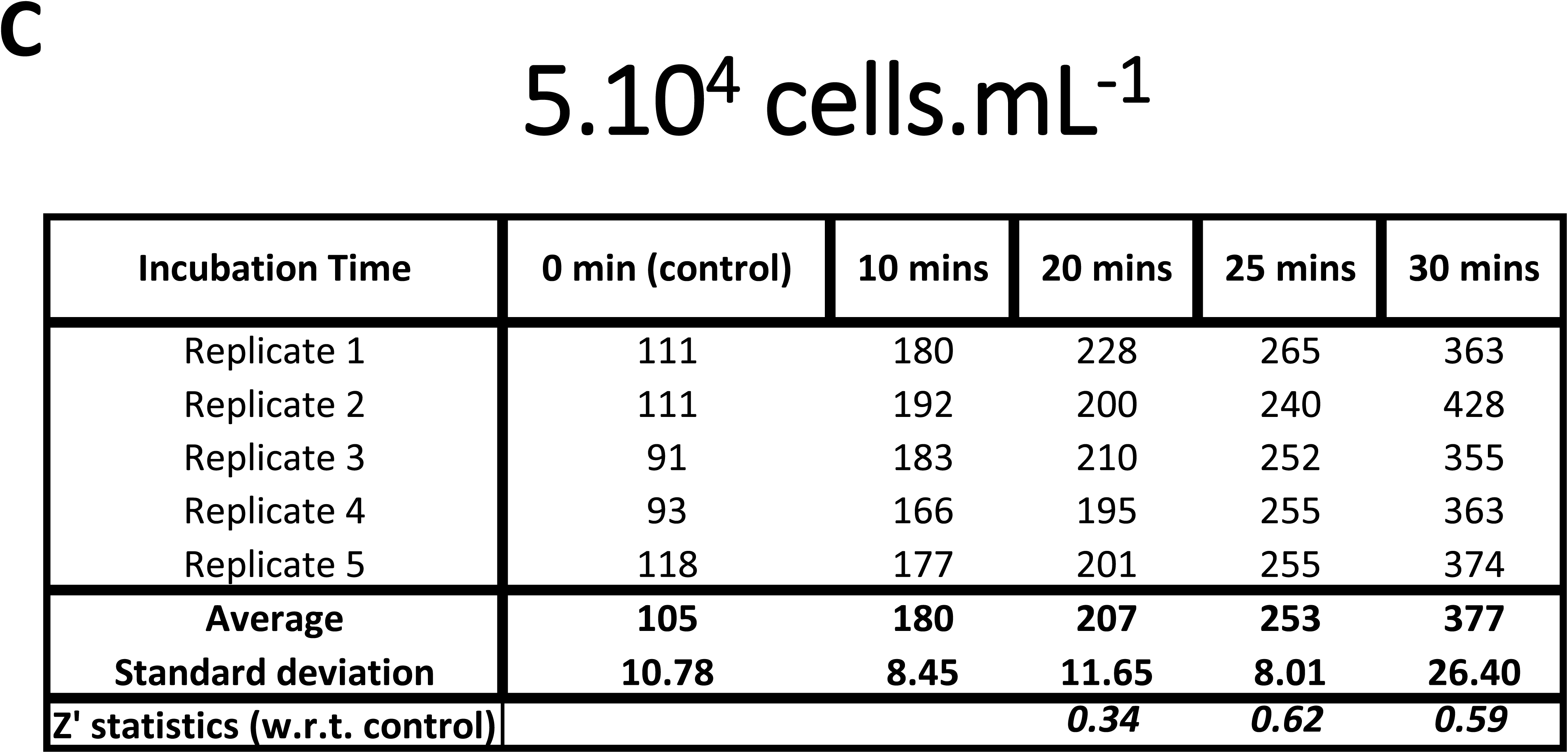
Changes in cell number during first 30 min following inoculation of cells from stationary phase into Terrific Broth. **A**. Typical cytograms. **B**. Reproducibility and Z’ statistics for *E. coli* growth at initial concentration of 10^5^ cells.mL^-1^. **C**. Reproducibility and Z’ statistics for *E. coli* inoculated at 5×10^5^ cells.mL^-1^.

### Flow cytometric assessment of antibiotic sensitivity

Figure 4 shows similar data for a resistant (Fig 4A,B) and a sensitive strain (Fig 4C,D) in the absence ((4A,C) and presence (Fig 4B,D) of the antibiotic ampicillin, applied at three times the known MIC (MIC = 32 mg.L^-1^) (http://www.eucast.org/fileadmin/src/media/PDFs/EUCAST_files/Breakpoint_tables/v_8.1_Breakpoint_Tables.pdf). It is clear that the susceptible strain differs (and thereby can be discriminated) from the resistant strain in at least three ways: (i) the kinetics of changes in cell numbers as judged by RL1 counts, (ii) the same as judged by forward (not shown) or side scatter, (iii) kinetic changes in the magnitude of the fluorescence.

**Figure 4.**
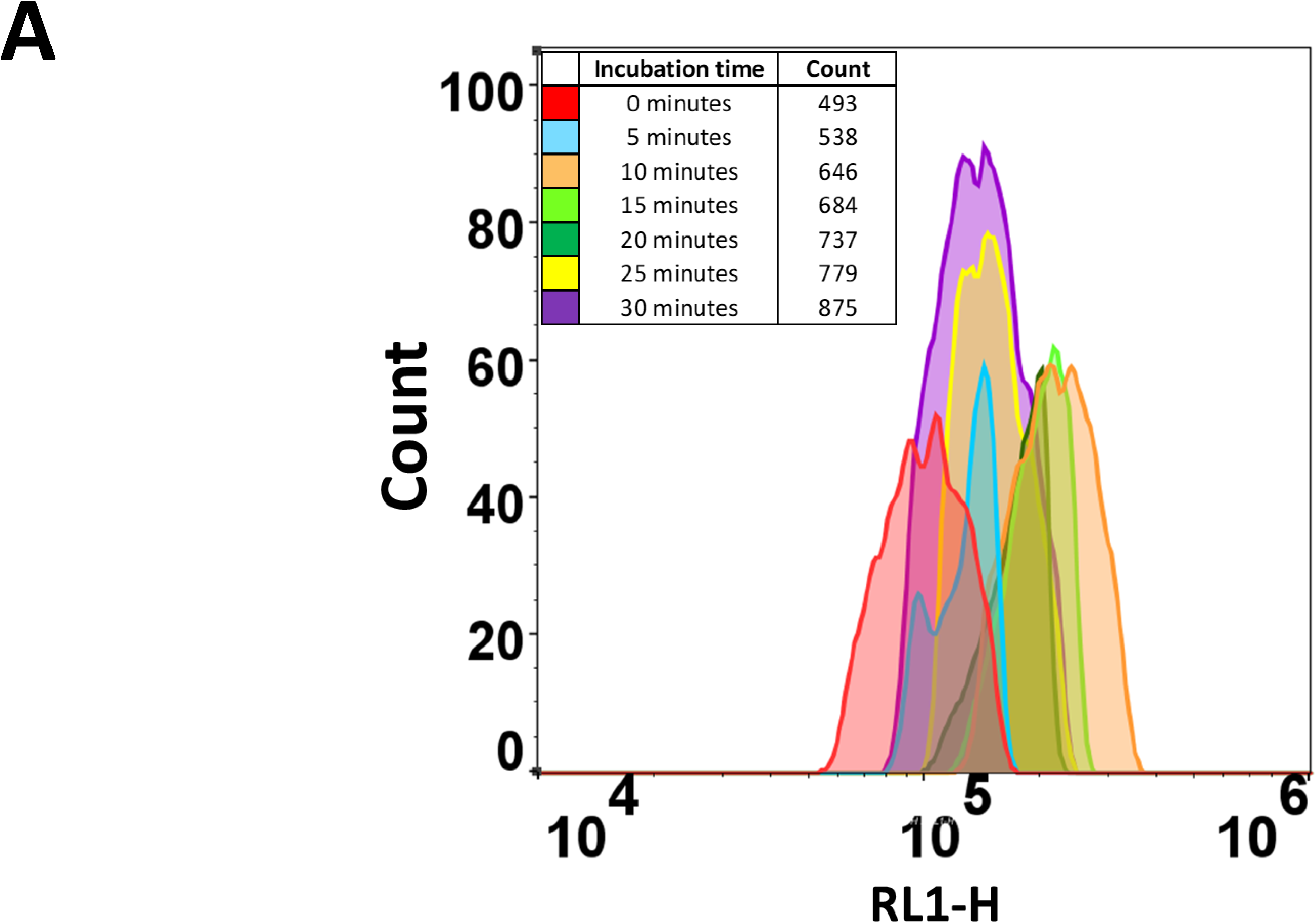
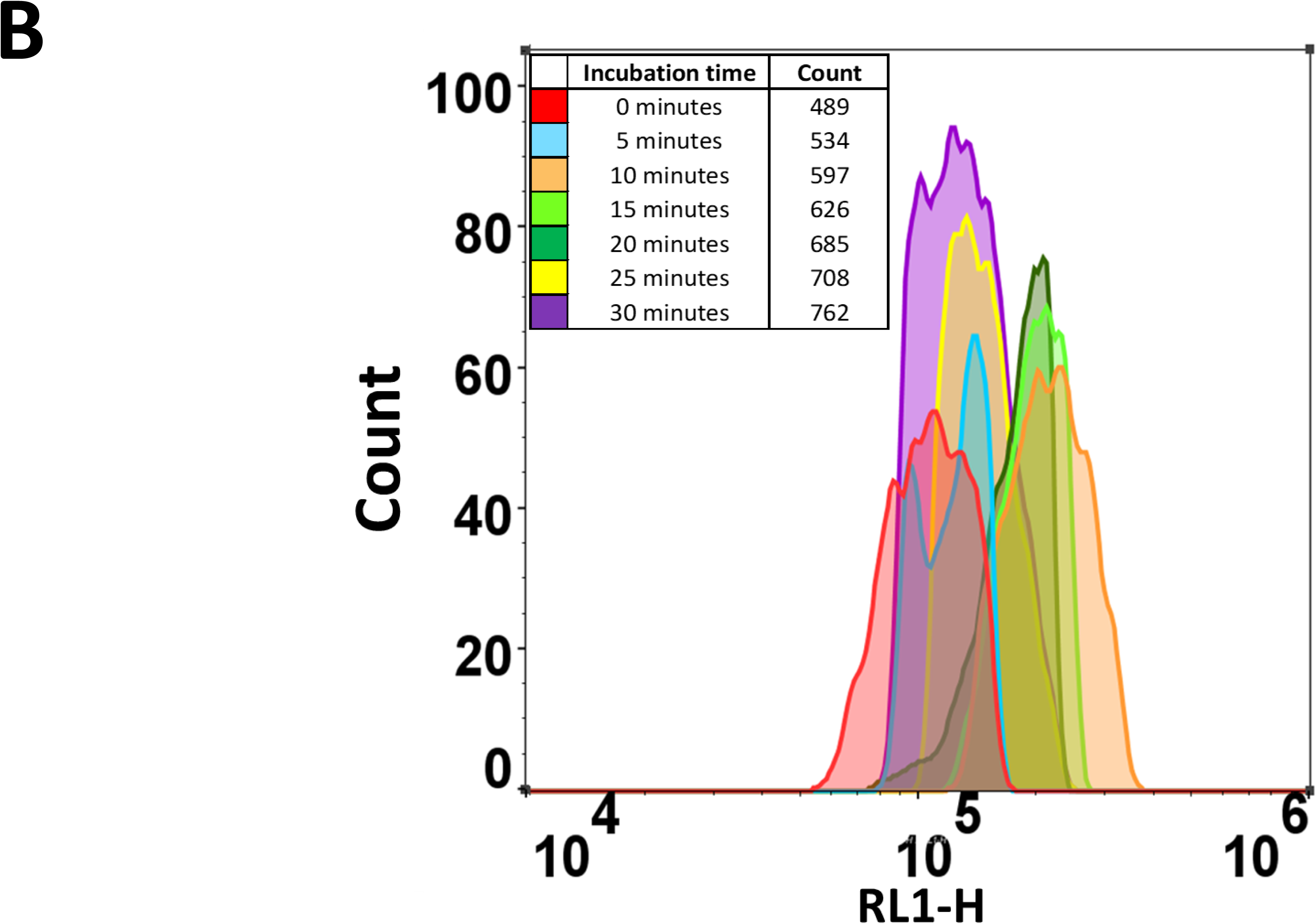
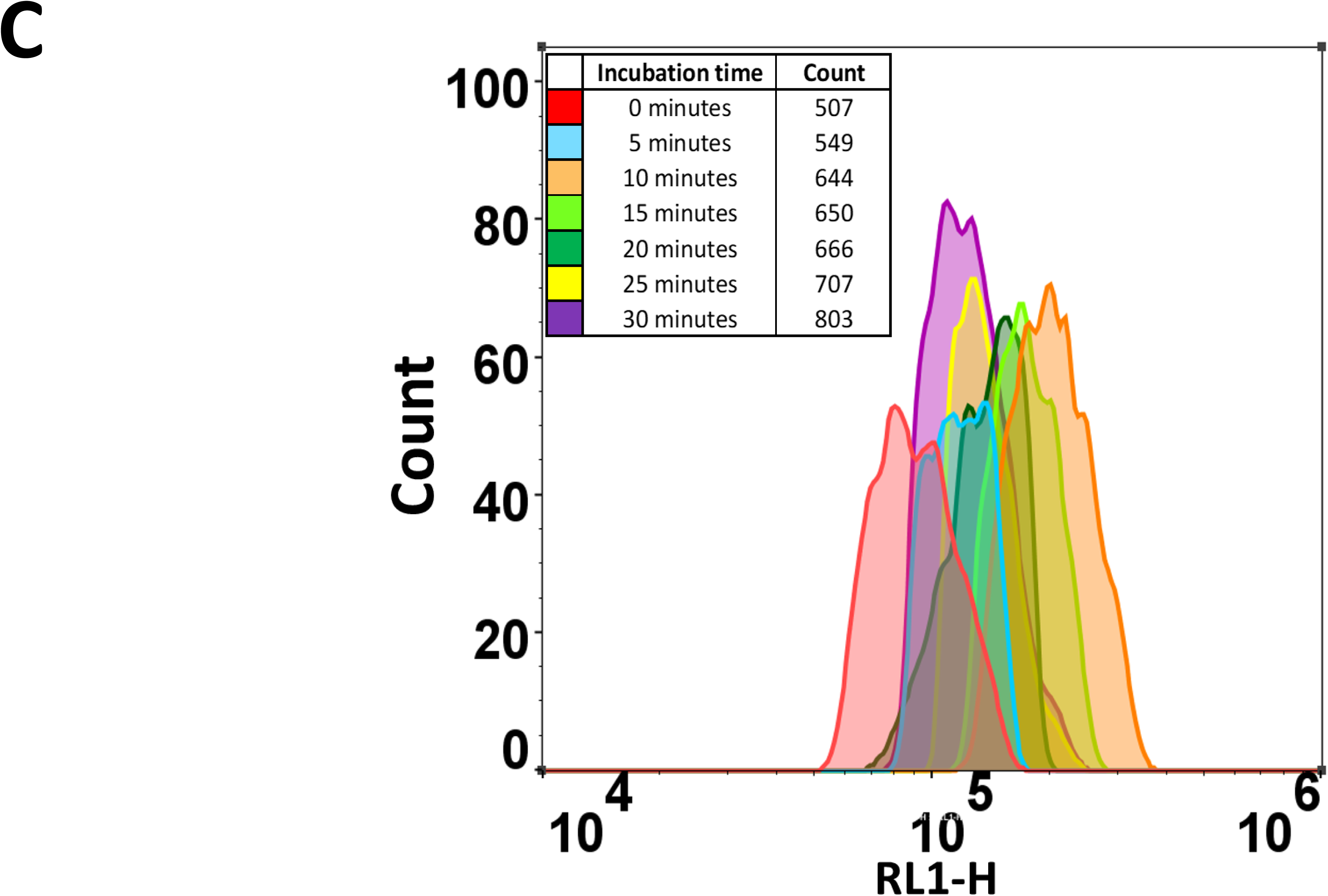
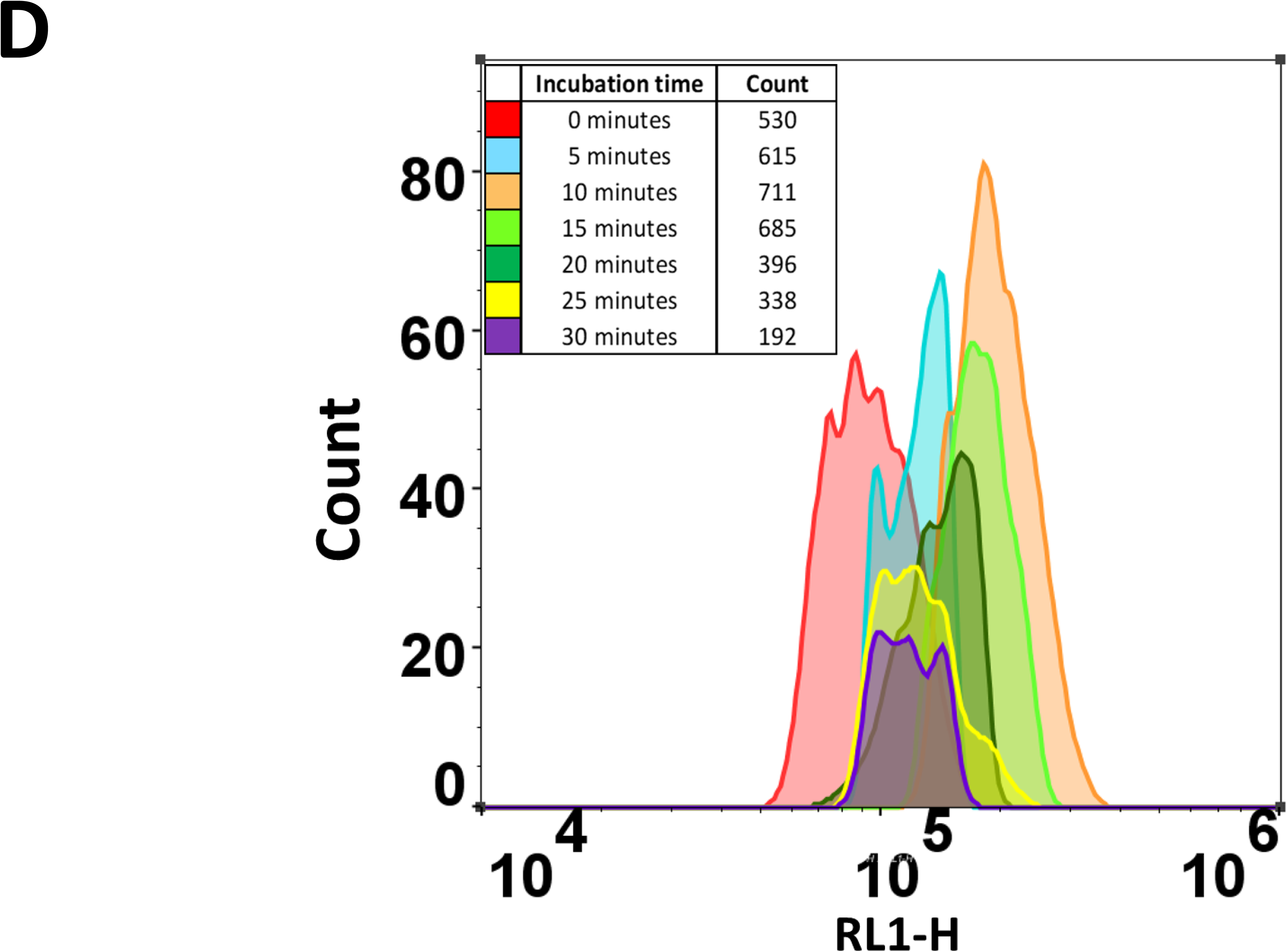
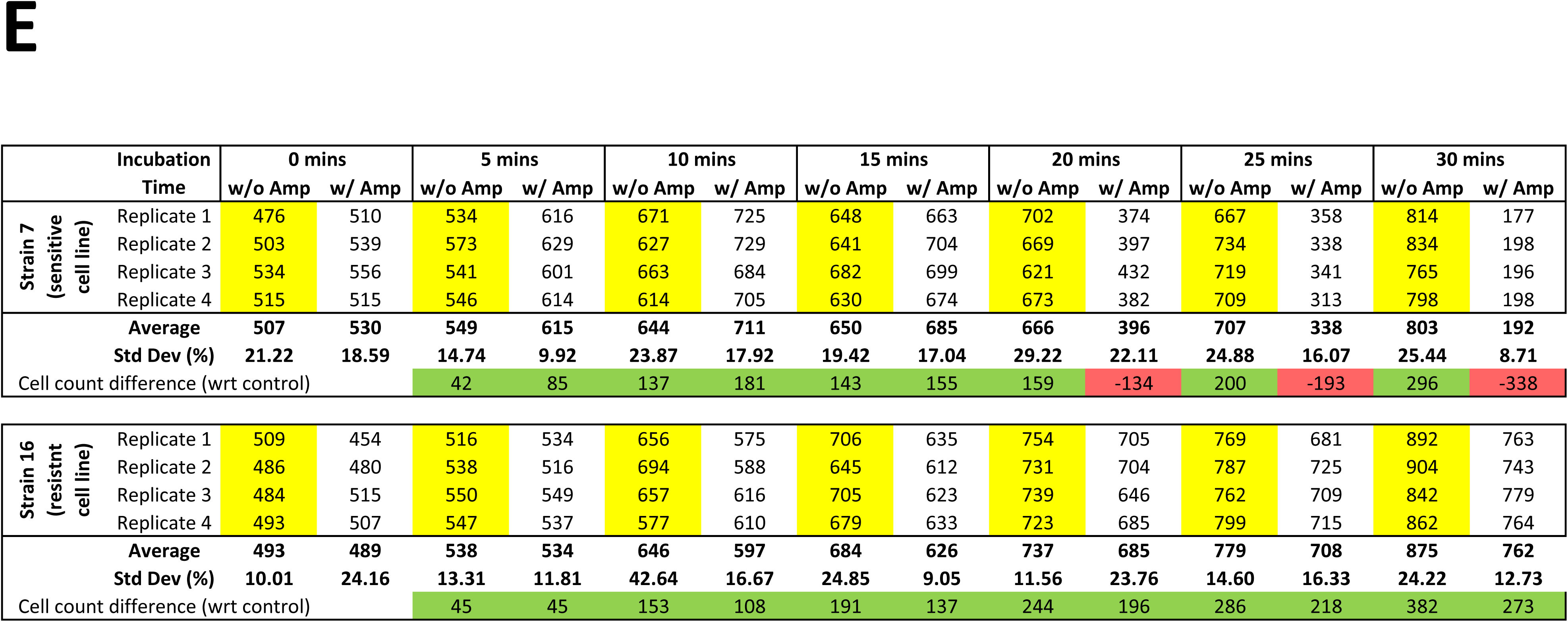
Effect of ampicillin (100 mg.L^-1^ concentration, 3 x MIC for sensitive strains) on the cytograms of *E. coli* inoculated from stationary phase into Terrific Broth. Ampillicin was either absent (**A,C**) or present (**B,D**) from resistant strain 16 (**A**,**B**) or sensitive strain 7 (**C**,**D**). **E.** Table showing the changes in the number of bacteria (with replicates) from sensitive (strain 7) and resistant strains (strain 16) when grown in the presence and absence of Ampicillin. Similar data were obtained using eight other macroscopically sensitive and resistant strains.

Since we had seen rapid changes in side scatter within 5 min (Figure 3) it was also of interest to study this as a means of detecting antibiotic sensitivity. Figure 5 shows that the changes in side scatter also differs noticeably between sensitive and resistant strains in 5-10 minutes, albeit that limited proliferation was taking place.

**Figure 5.**
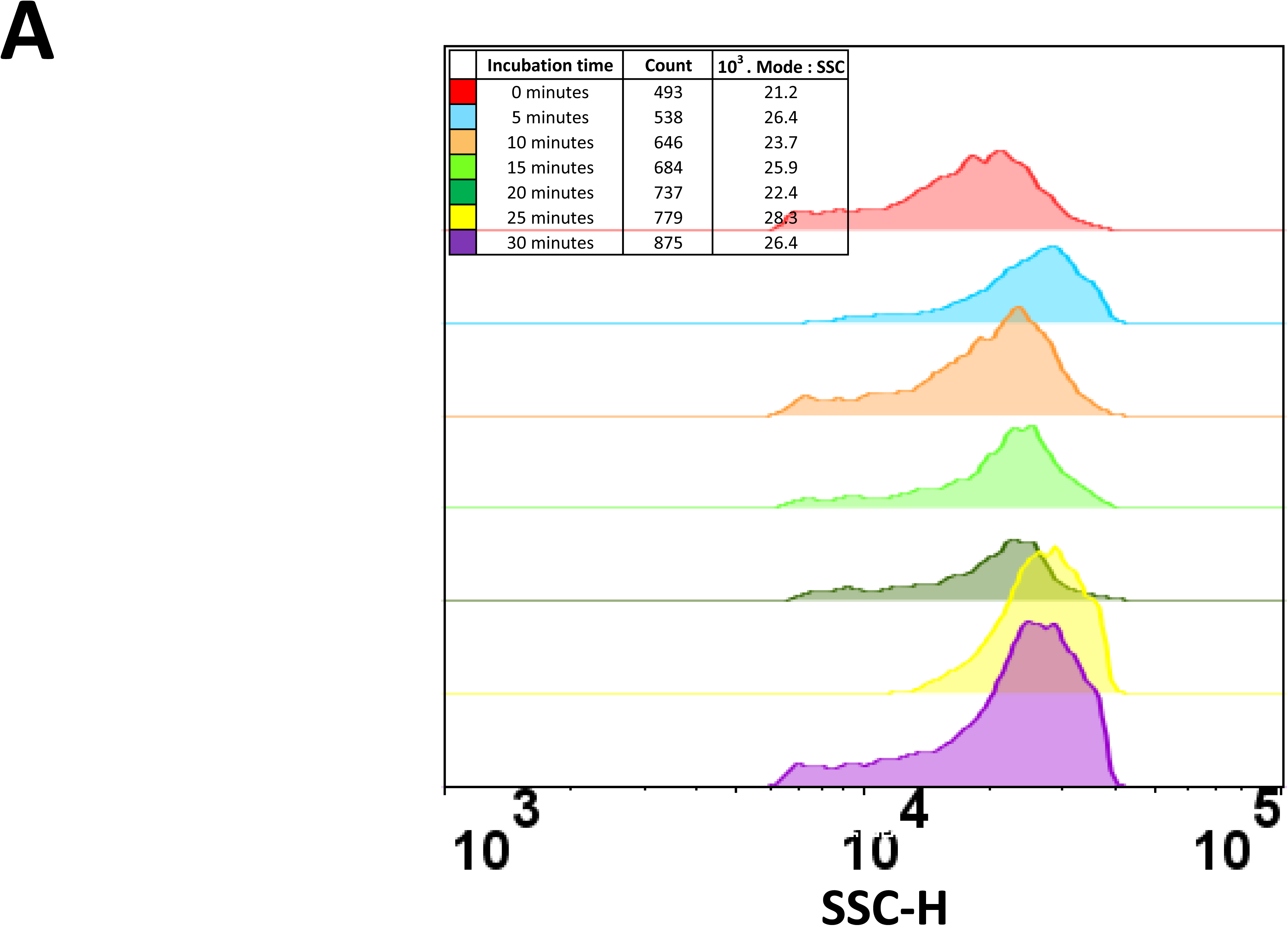
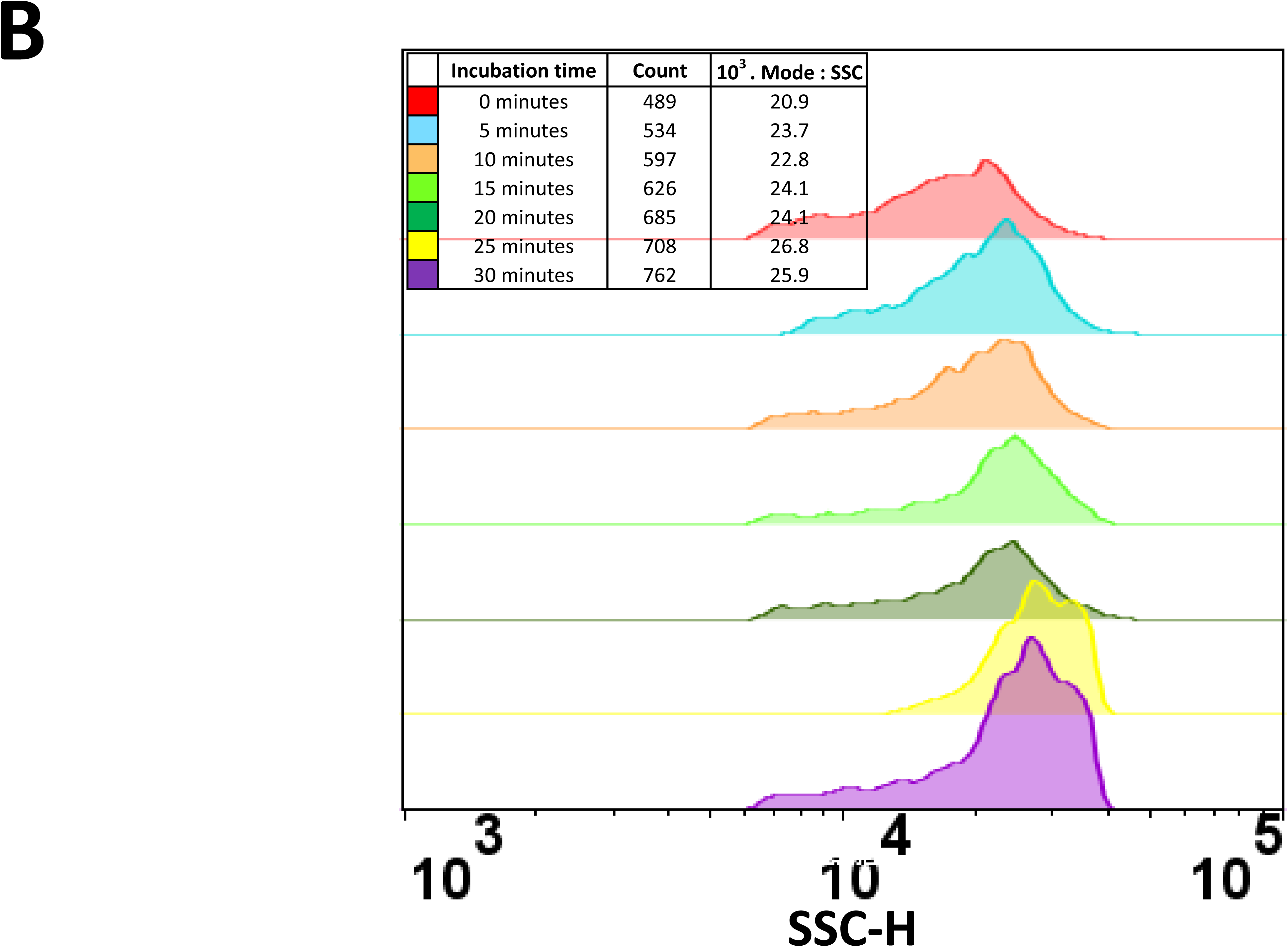
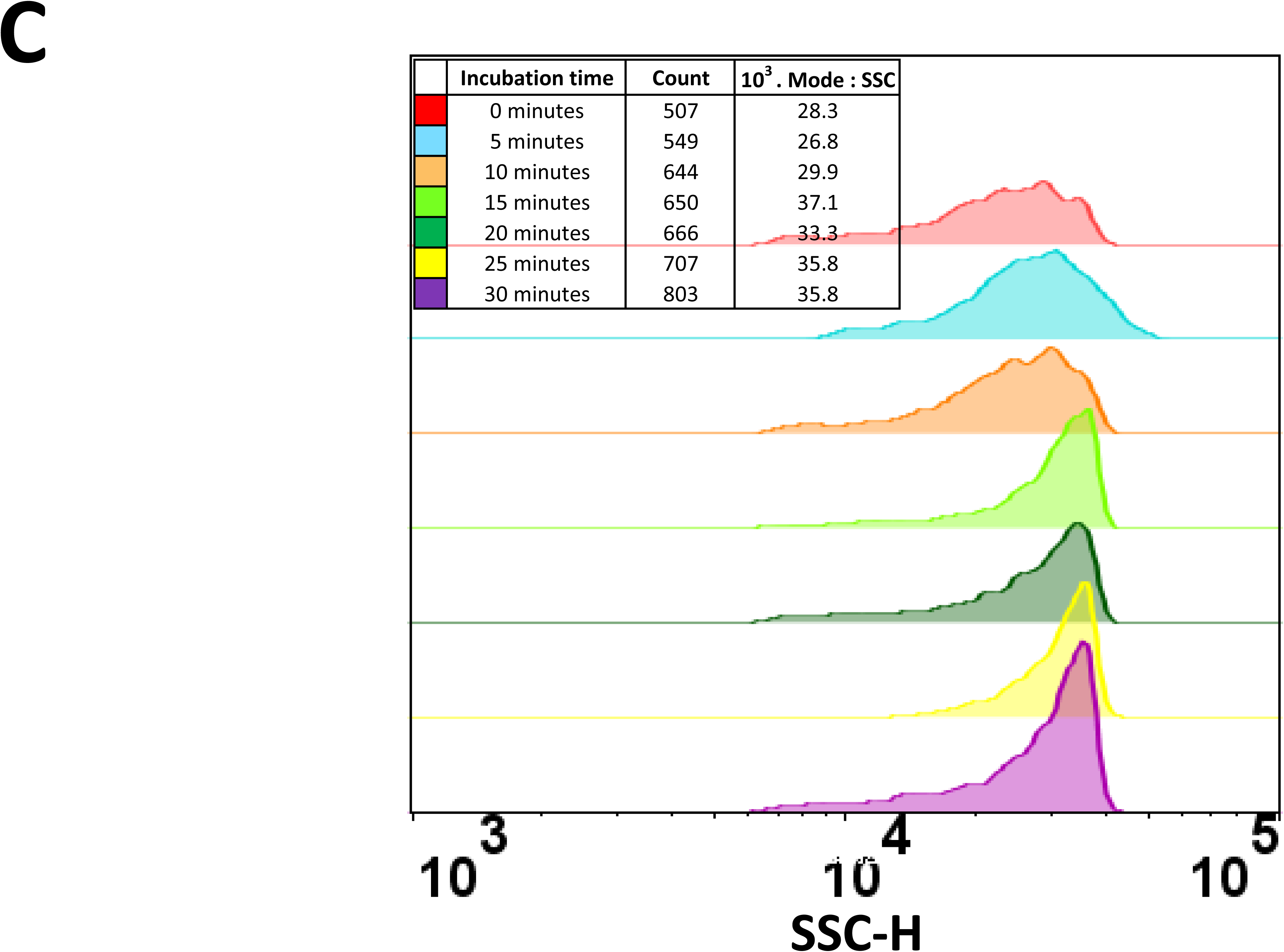
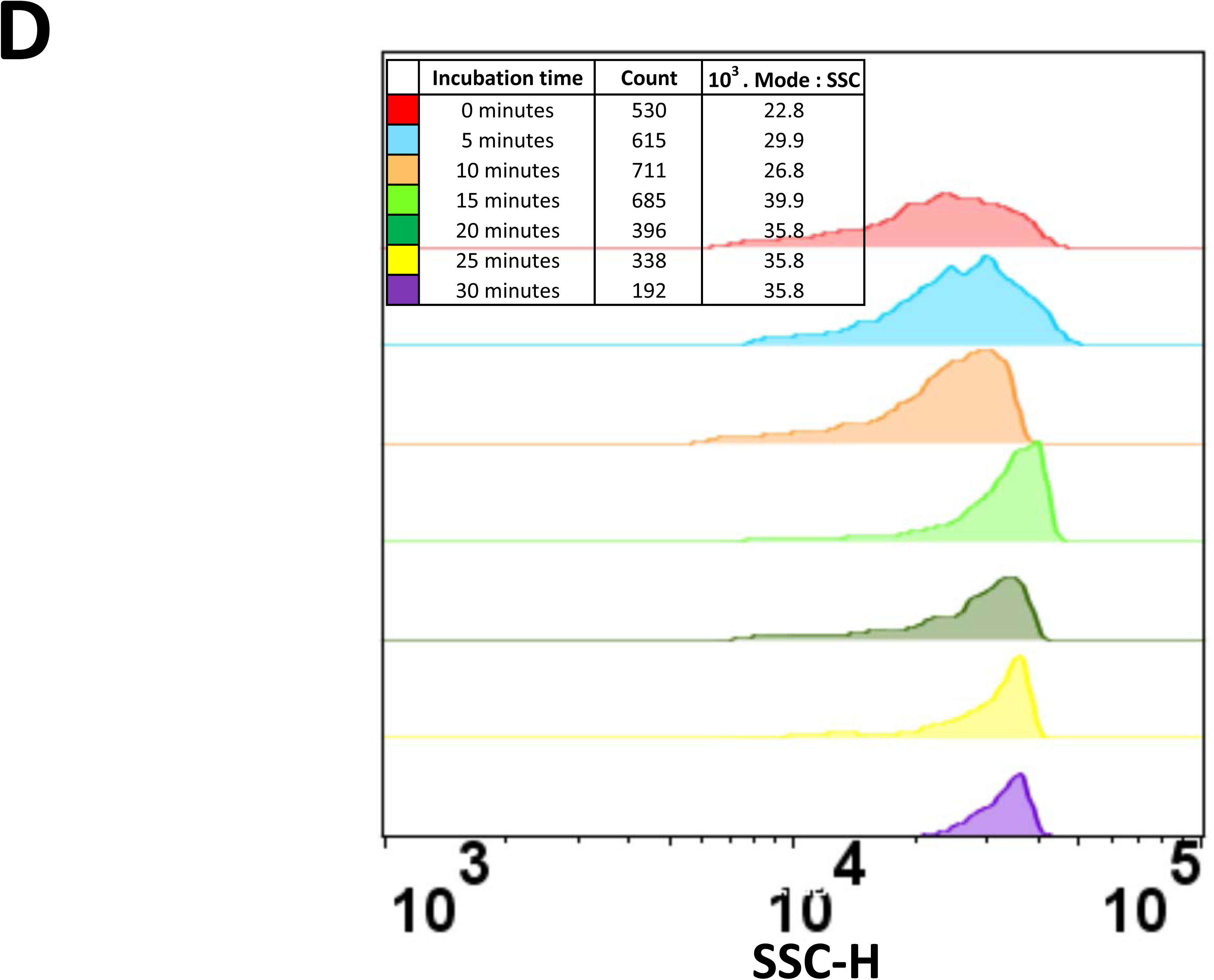
Side scatter histograms of the experiment mentioned in Figure 4. Ampillicin was either absent (**A,C**) or present (**B,D**) from resistant strain 16 (**A**,**B**) or sensitive strain 7 (**C**,**D**).

Of course different antibiotics have different modes of action [118, 119], and the optimal readout needs to reflect this. Thus, nitrofurantoin is widely prescribed for UTIs and its effects on our standard laboratory system are shown in Fig 6A,B (cytograms of side scatter and RL1, respectively). The effects on cell proliferation of nitrofurantoin and several other antibiotics are given in Fig 6C. Note that the initial and later cell numbers for nitrofurantoin appear lower because this antibiotic absorbs light at the excitation wavelength (its peak is at 620 nm). Both the bacteriostatic (trimethoprim) and bactericidal (ampicillin, ciprofloxacin, nitrofurantoin) antibiotics can be seen to work effectively on this sensitive strain.

**Figure 6.**
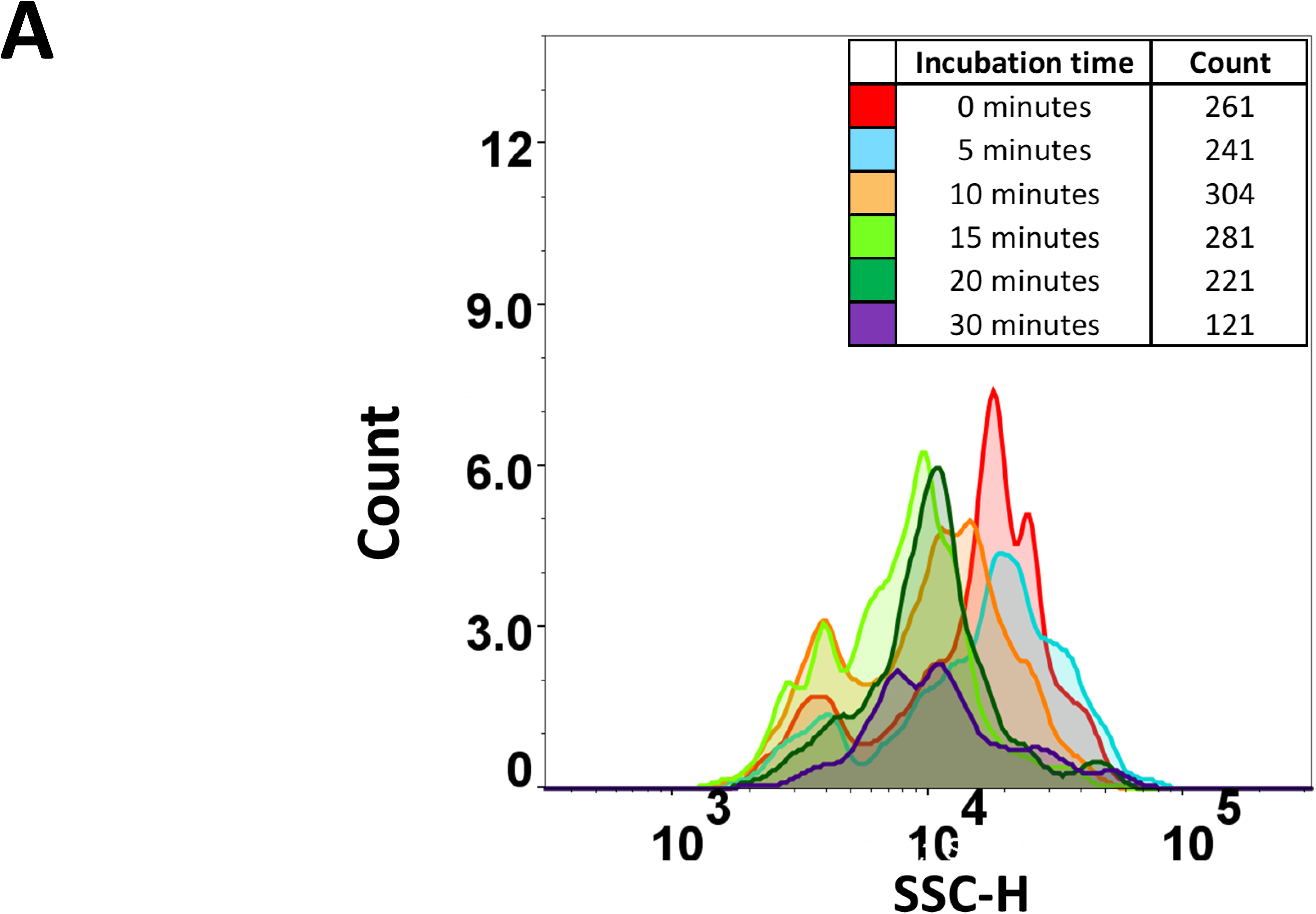
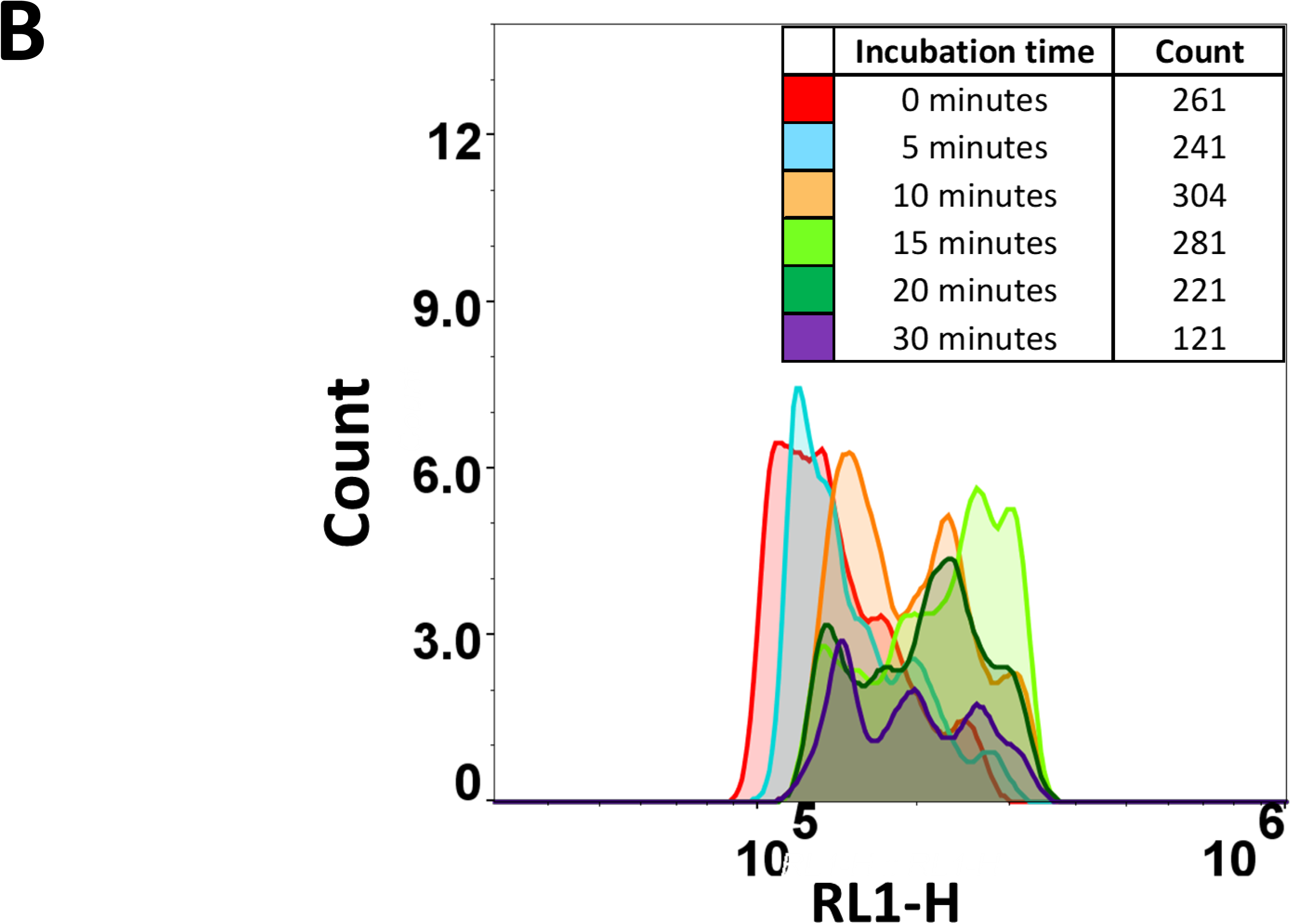
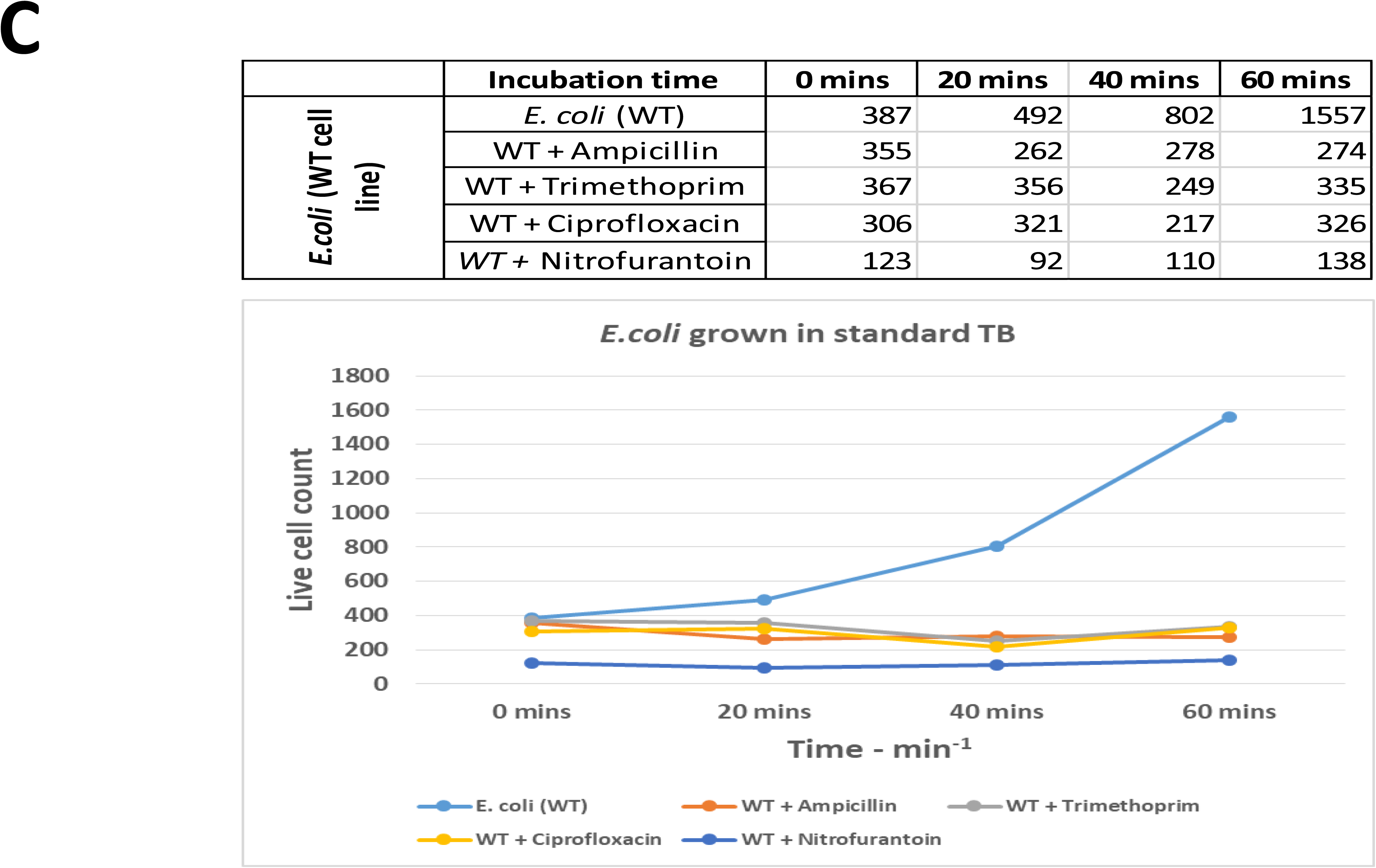
Effect of nitrofurantoin at 3x nominal MIC on the growth and flow cytometric behaviour of a sensitive strain of E. coli. **A,B** for nitrofurantoin, cytograms of (**A**) side scatter, (**B**) RL1 fluorescence. Experiments were performed precisely as shown in the legend to Fig 3. (**C**) Ability of flow cytometric particle counting (gated as in Fig 2) to determine the sensitivity of *E. coli* MG1655 to four different antibiotics in 20 mins.

### Flow cytometric assessment of DNA distributions

Another important strategy for detecting bacteria uses their DNA (e.g. [120-122]). Thus, another high-level guide to the physiology of *E. coli* cells and cultures is the flow cytometrically observable distribution of DNA therein, as this can vary widely as a function of growth substrate, temperature, and during the cell cycle [39-43]. Specifically, the solution to the problem that DNA replication rates are fixed while growth rates can both vary and exceed them is to allow multiple replication forks in a given cell [123]. To this end, we compared the DNA distributions of our cultures under various conditions. Fig 7A shows both stationary phase and exponentially growing cells stained with a mithramycin-ethidium bromide cocktail as per the protocol of Skarstad and colleagues given in Materials and Methods. As they have previously observed [40, 47], (very slowly growing or) stationary phase cells display either one or two chromosome complements, while those growing exponentially in lysogeny broth (LB medium) can have as many as eight or more chromosomes. This is entirely consistent with the basic and classical Cooper-Helmstetter model [123] and more modern refinements [124-127]. To this end, Fig 7B shows changes in the DNA distribution of cells taken from a similar regrowth experiment to that in Fig 2. It is evident that both the one- and two-chromosome-containing cells from the stationary phase initiate increases in their DNA content on the same kinds of timescale as may be observed from both direct cell counting (proliferation) and carbocyanine fluorescence, with the initially bimodal DNA distribution morphing into a more monomodal one. This implies that the initial increase in cell numbers over 15 min or so involves cells that were about to divide actually dividing, and provides another useful metric of cellular (cell cycling) activity, albeit one that requires sampling as the cells must be permeabilised, at least for this protocol.

**Figure 7.**
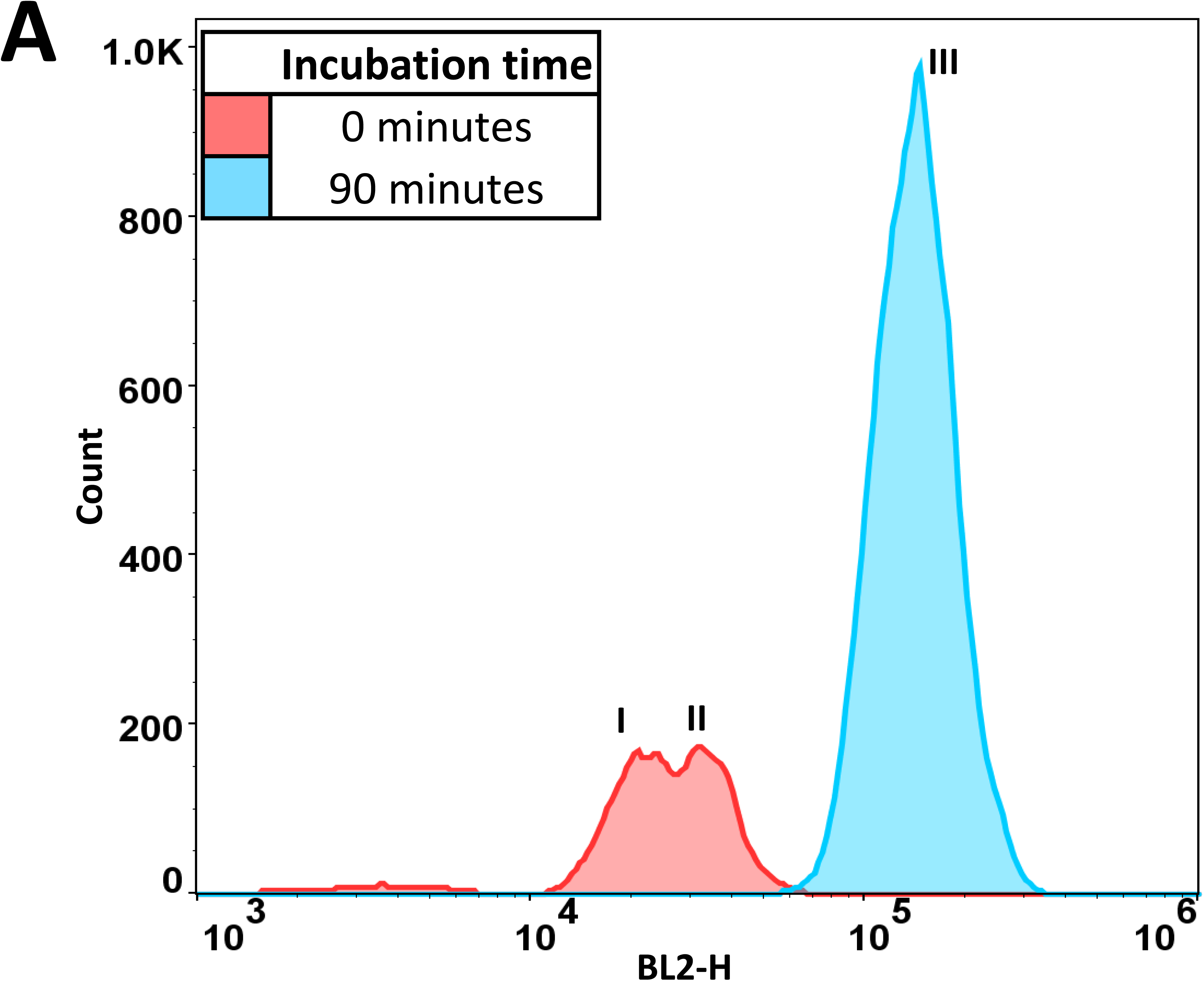
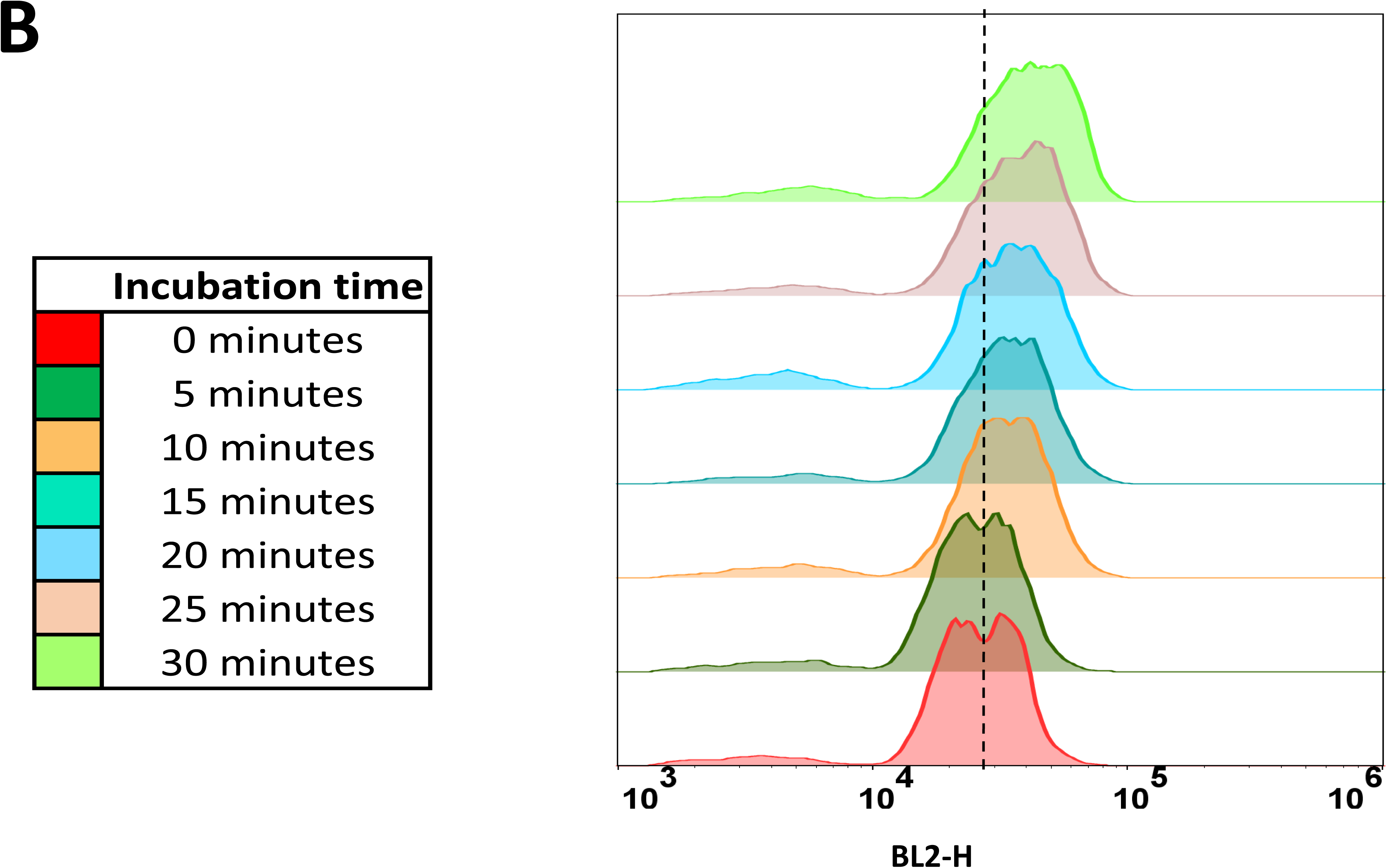
DNA distributions in different populations. **(A).** DNA distributions in stationary phase (red) and exponentially growing cells (blue). The overlay histogram shows data from *E. coli* samples that were fixed with 70% ice cold ethanol and then stained using Mithramycin and Ethidium Bromide as described in Materials and Methods. The relative intensity of the BL2 channel fluorescence (488nm excitation, 572±14 nm emission) shows the amount of chromosomes in the cells. The points I, II and III represent one, two and eight chromosome equivalents, respectively. The peak values of BL2 fluorescence for the points are I (2.03.10^4^), II (3.90.10^4^) and III (1.53.10^5^). The cells in stationary phase (2-4h) were taken and fixed immediately while the exponentially growing cells were incubated for 90 minutes at 37°C before fixing the cells. **(B)**. Changes in DNA distribution in *E. coli* cells following inoculation from a stationary phase into Terrific Broth every 5 min until 30 min. Experiments were otherwise performed exactly as described in the legend to in Figure 2.except that (to avoid spectral interference) carbocyanine was not present.

## Flow cytometric analysis of UTI samples

Finally, we wished to determine whether this method, as developed in laboratory cultures, could be applied to candidate UTI specimens ‘as received’ in a doctor’s surgery. To this end, we analysed 23 samples of (initially) unknown properties, of which six were in fact positive. Each of these was found to be positive using our methods, and with the antibiotic sensitivities given in Table 1. Typical cytograms for sensitive and resistant strains are given in Fig 8. The positive cultures were speciated centrally, and in each case the organism was found to be *E. coli*.

**Table 1.** Antibiotic sensitivity profile for the six positive samples (taken to be ≥10^5^.mL^-1^) obtained from the Firsway clinic. Two separate samples were from 25/05/18.

|  | Antibiotic sensitivity (R- resistant; S- sensitive) |  |  |  |
| --- | --- | --- | --- | --- |
| Sample Date | Ampicillin | Trimethoprim | Ciprofloxacin | Nitrofurantoin |
| 25/05/2018 | R | R | S | S |
| 25/05/2018 | S | S | S | S |
| 06/06/2018 | R | S | S | S |
| 16/07/2018 | R | S | S | S |
| 18/07/2018 | R | S | R | R |
| 20/07/2018 | S | R | S | S |

**Figure 8.**
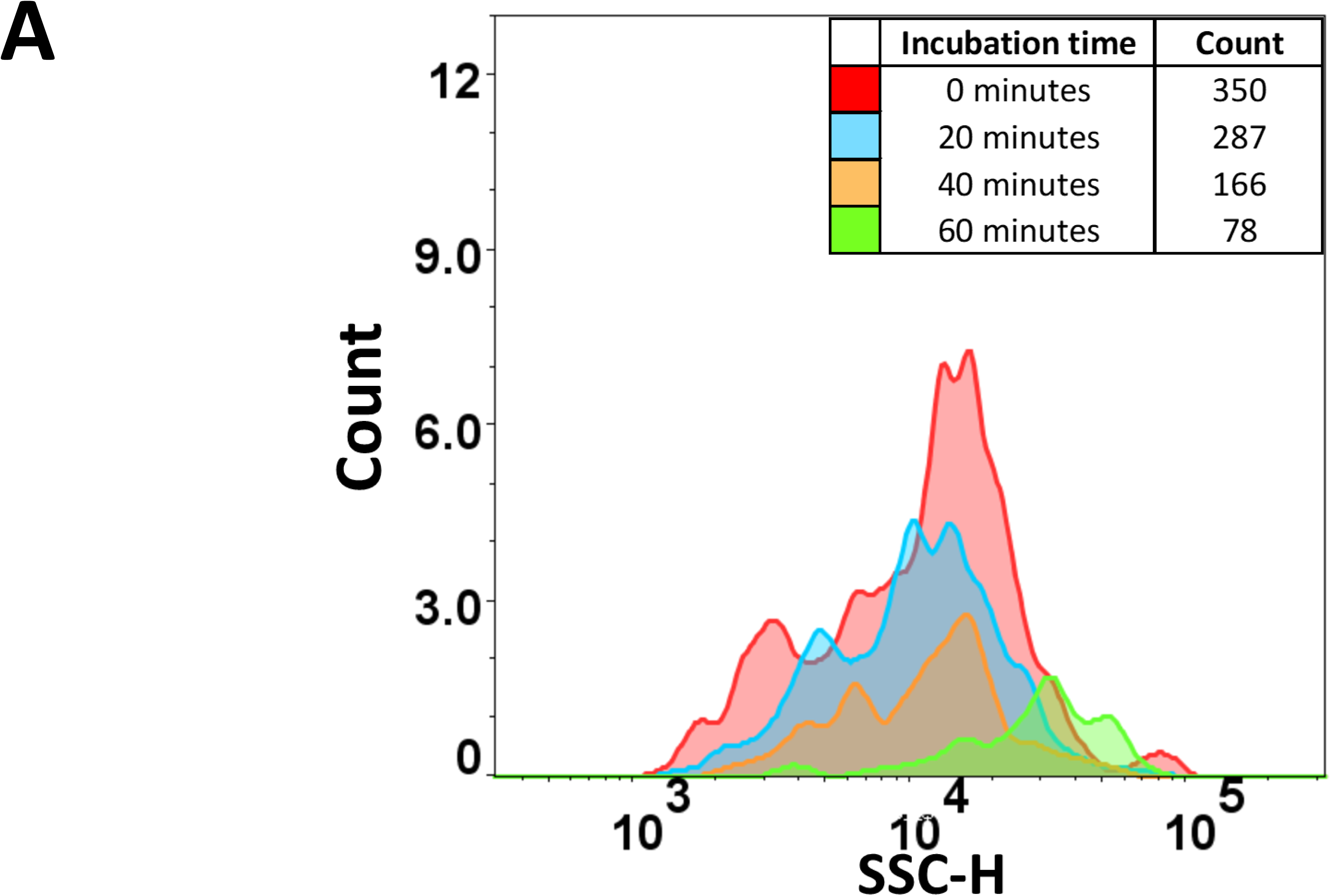
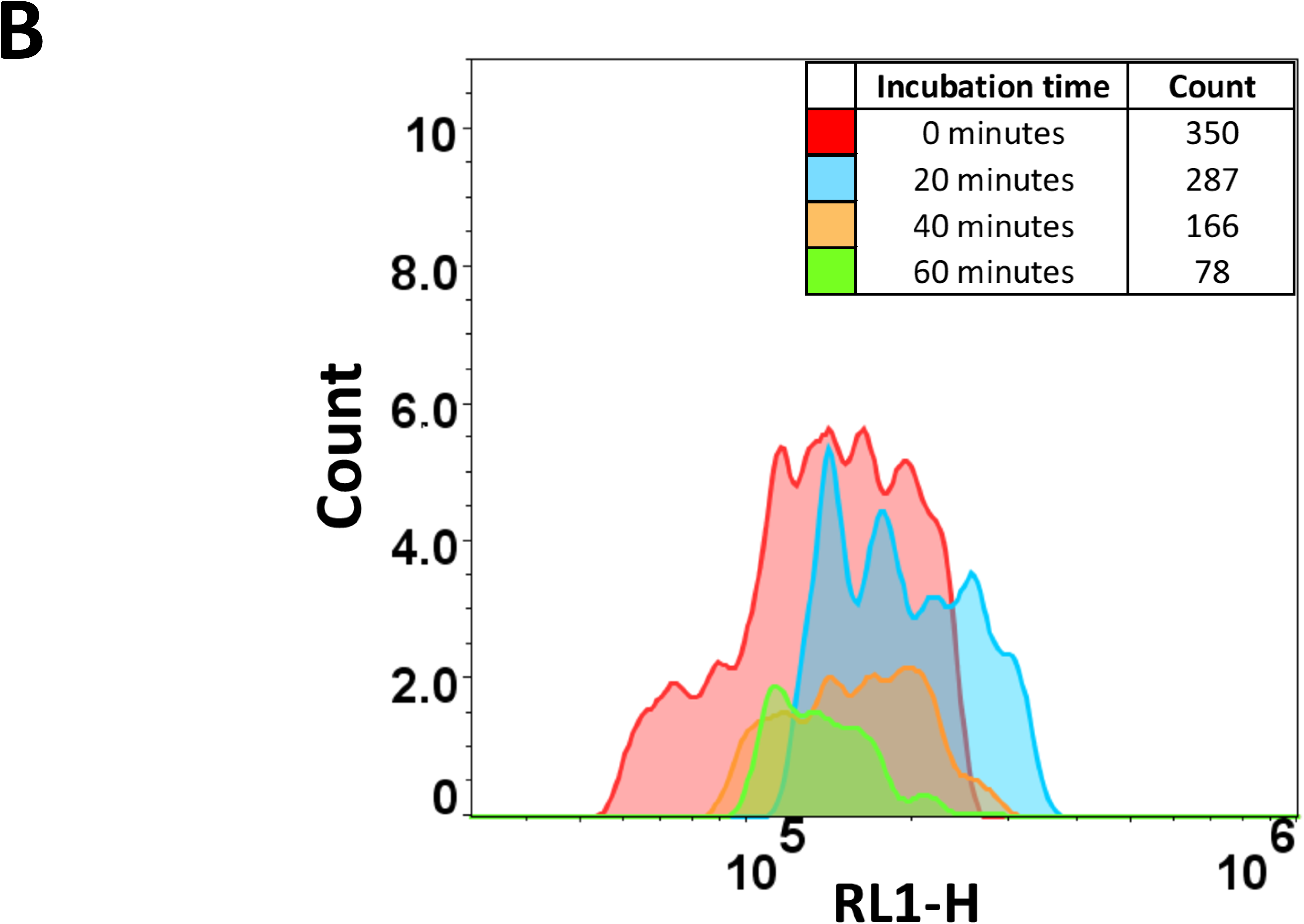
Cytograms of a sensitive UTI strain treated with nitrofurantoin. UTI samples (in this case containing ~ 10^6^ cells.mL^-1^) were taken directly from storage, diluted tenfold into 37°C Terrific Broth including 3 µM diS-C3(5) and nitrofurantoin at a nominal 3x MIC, and measured flow cytometrically as described in the legend to Fig 2. (**A**) side scatter, (**B**) red fluorescence.

## Discussion and conclusions

It is often considered that the ‘lag’ phase of bacterial growth is one in which very little is happening, and that what is happening is happening quite slowly. This notion probably stems from the fact that changes in OD observable by the naked eye in laboratory cultures [77] are indeed quite sluggish. However, the very few papers that have studied this in any detail [63, 84, 85, 95, 97, 98, 102, 103] have found that changes in expression profiles (albeit mainly measured at a bulk level) actually occur on a very rapid timescale indeed, possibly in 4 minutes or less following reinoculation into a rich growth medium. For antibiotics to have an observable, and in terms of sensitivity to them a <u>differentially</u> observable, effect on cells, the cells need to be in a replicative state. This might be thought to preclude any such observations in the lag phase, but what is clear from the present observations is that cells can re-initiate or continue their cell cycles very rapidly, such that observable proliferation can occur in as little as 15-20 min after reinoculation of starved, stationary phase cells into rich medium. Consequently it is not necessary to wait for a full period of ‘lag-plus-first-division time’ [63], which can be well over one hour [115, 116]. The rapid proliferation that we describe could be observed by light scattering, by cell counting, by carbocyanine fluorescence (membrane energisation), and by changes in the magnitude and distribution of DNA in the population. This has allowed us to determine, using any or all of these phenotypic assays, antibiotic susceptibility at a phenotypic level in what would appear to be a record time. Pin and Baranyi [115, 116] observed a more stochastic and somewhat slower process than that which we observed here, but in their case they were measuring CFU only, and the inoculation was into the less rich LB, while we used Terrific Broth. Indeed, the exit from lag phase can be very heterogeneous when organisms are measured individually [63, 128-130].

While we did not study this at the level of the transcriptome here, the dynamics of the physiological changes observed during the early lag and regrowth phases as observed by the uptake of the carbocyanine dye are of interest. Classically, its uptake has been considered to reflect a transmembrane potential difference (negative inside) (e.g. [78, 79, 131-134], but cf. [135]) based on bilayer-mediated equilibration according to the Nernst equation [136]. However, we recognise that such cyanine dyes, much as ethidium bromide [120] and other xenobiotics [137, 138], are likely to be both influx and efflux substrates for various transporters [139], so such an interpretation should be treated with some caution.

As to future work, the same strategy may usefully be applied to other cells (including pathogens in both urine and more difficult matrices), other antibiotics and other stains. However, the present work provides a very useful springboard for these by showing that one may indeed expect to be able to determine antibiotic susceptibility in a phenotypic assay in under 20 minutes. This could be a very useful attribute in the fight against anti-microbial resistance.

## Acknowledgments

We thank the UK Biotechnology and Biological Sciences Research Council (BBSRC) for financial support (grant BB/R000093/1), Dr Howbeer Muhamadali for access to various strains, and Prof Roy Goodacre for useful discussions. Stuart Logan and Sun Yung provided invaluable advice and assistance during the commissioning of the Intellicyt instrument.

## Abbreviations

DiBAC4(3): bis-(1,3-dibutyl-barbituric acid) trimethine oxonol
di-S-C3(5): 3,3′-dipropylthiadicarbocyanine iodide

